# Human-targeted drugs extensively reshape regulatory networks controlling virulence and antibiotic resistance in *Salmonella* Typhimurium

**DOI:** 10.64898/2026.09.30.755304

**Authors:** Martin K. Amstalden, Roberto Olayo-Alarcón, Susanne Brenzinger, Lorine Debande, Annamaria Zannoni, Elisabetta Fiore, Abdelhakim Boudrioua, Azam Rashidian, Thales Kronenberger, Samuel Wagner, Cynthia M. Sharma, Christian L. Müller, Ana Rita Brochado

**Author notes:** University of Luebeck, Institute of Medical Microbiology, University Hospital Schleswig-Holstein, Campus Luebeck, Germany (Abdelhakim Boudrioua).

## Abstract

Human-targeted drugs are increasingly recognized to affect bacterial physiology, yet their impact beyond growth inhibition, and the underlying molecular mechanisms, remain poorly understood. Here, we systematically mapped how chemically diverse host-associated compounds influence virulence and intrinsic antibiotic resistance in *Salmonella enterica* serovar Typhimurium. Using a high-throughput reporter approach, we quantified transcriptional response of 26 key stress promoters in *Salmonella* against 2,415 compounds, including FDA-approved drugs, food-derived molecules, and human metabolites. We found that non-antibiotic compounds induce transcriptional responses distinct from those elicited by antibiotics, uncovering a regulatory shift from general stress responses to compound-specific regulation driven by ligand-responsive MarR- and TetR-family regulators. In particular, more than 200 non-antibiotic drugs modulate virulence or intrinsic resistance pathways through the central regulators SlyA or RamR, respectively. By integrating our large-scale dataset with machine learning, we further defined chemical features associated with *ramA* transcriptional activation in *Salmonella*, and predicted ∼200 additional candidate inducers of intrinsic antibiotic resistance in this pathogen. Together, our results uncovered a large and previously unrecognized chemical landscape that reshapes bacterial stress responses through ligand-responsive regulatory mechanisms, with implications for how host-associated compounds influence pathogen virulence and antibiotic resistance.

**Highlights:**

- Mapping the transcriptional response of key *Salmonella* stress pathways across a large chemical space shows that antibiotics and non-antibiotic compounds elicit distinct stress responses in *Salmonella*
- The activity of the virulence regulator SlyA is modulated by non-antibiotic compounds
- SlyA-inhibitors increase sensitivity to polymyxin B and decrease *Salmonella* virulence in cell culture
- Human-targeted drugs increase intrinsic antibiotic resistance via RamR-RamA in *Salmonella*
- Machine learning approach enables large-scale, out-of-sample prediction of human-targeted drugs inducing *ramA* expression in *Salmonella*

## Introduction

It is well acknowledged that human-targeted drugs can profoundly impact bacterial growth and the composition of microbial communities^1,2^. At the same time, emerging evidence indicates that human-targeted drugs can also modulate clinically relevant phenotypes beyond growth, including antibiotic resistance, antibiotic interactions, virulence, and colonization resistance^3–7^. However, the molecular mechanisms underlying these effects, as well as the broader chemical landscape of compounds that influence bacterial phenotypes beyond growth, remain largely unexplored. This gap in knowledge is particularly important given the widespread and often concurrent use of these drugs. Here we investigate the broad impact of host-related small-molecules, including human-targeted drugs and food-homology compounds, on the regulation of intrinsic antibiotic resistance and virulence of the model pathogen *Salmonella enterica* serovar Typhimurium (hereafter *Salmonella*).

*Salmonella* is a Gram-negative bacterium, very well studied for virulence and pathogenicity, with notable intrinsic antibiotic resistance, primarily mediated by proficient efflux pumps^8,9^. Regulation of virulence and antibiotic resistance is embedded in a complex stress response network, which *Salmonella* uses to adapt to changing environments, e.g., during infection, nutrient change and antibiotic treatment^10–17^. Stress adaptation by *Salmonella* involves a large and diverse repertoire of so-called “stress response pathways” that tightly regulate and reshape gene expression. The extent of expression rewiring can vary from a handful of specific genes to 10-20% of the genome and is highly dependent on the stress condition^18–20^. Stress responses can be mediated by a variety of global regulators such as the sigma factors RpoS, RpoE, the transcription factor OxyR, and the PhoPQ and CpxAR two-component systems. These respond to perturbations in cellular homeostasis, including oxidative stress, membrane damage, or nutrient limitation. For example, PhoPQ senses low Mg²⁺ levels^21^, as well as host-relevant signals such as acidic pH and antimicrobial peptides, to orchestrate a transcriptional response involving several virulence genes^10^. In parallel, bacteria can also directly sense small molecules through ligand-responsive transcription factors in the cytosol, notably members of the MarR- and TetR-families^22,23^. These regulators typically repress genes involved in multidrug resistance, metabolism, and virulence, and relieve repression upon ligand binding^20,23–26^.

In *Salmonella*, the TetR-family regulator RamR controls expression of *ramA*, an activator of the AcrAB efflux pump, which is a major resistance determinant in this bacterium^20,26–28^. Additionally, increased *ramA* expression via *ramR* mutations is frequently observed in resistant clinical isolates and associated with multi-drug resistance in *Salmonella*^26,29^. A number of compounds - including indole, bile acids, antibiotics, and a few human-targeted drugs - induce *ramA* expression^27,30,31^. For a limited subset of them, ligand-binding to RamR has been shown to trigger *ramA* overexpression^32,33^. Similarly, MarR-family regulators such as MarR, EmrR, and SlyA respond to small molecules, particularly aromatic carboxylates^34,35^. Among these three regulators, SlyA plays a central role in *Salmonella* virulence by counteracting H-NS-mediated gene silencing of pathogenicity genes^36,37^, while MarR and EmrR are most often associated with multidrug resistance through control of expression of efflux pumps^9,34,38,39^. Despite extensive characterization, getting a holistic view of how these pathways are engaged across diverse chemical environments remains very challenging, mostly due to pathway crosstalk, redundancy, and the largely unexplored range of activating signals. In particular, which stress pathways are activated in *Salmonella* in response to human-targeted drugs, and if and how this culminates in change in virulence and intrinsic resistance, remains largely unknown.

High-throughput chemical genomics and molecular profiling have become powerful approaches for systematically dissecting the molecular mechanisms that govern bacterial responses to environmental and chemical perturbations^38,40–43^. Here, we apply a high-throughput approach to systematically map the transcriptional response of selected antibiotic resistance- and virulence-associated genes to 2,415 compounds, including FDA-approved drugs, food-related molecules, and human metabolites, in *Salmonella* (Fig. 1a). In addition, we included representative reporters of canonical pathways, including oxidative, envelope, and RpoS-mediated stress response, to capture broader compound effects on *Salmonella* physiology. In total, we mapped a comprehensive network of 62,790 compound–promoter pairs, and show that antibiotics and non-antibiotics induce fundamentally distinct transcriptional responses. In addition, we reveal extensive regulation of drug transport and virulence in *Salmonella* by human-targeted drugs through the ligand-binding regulators RamR and SlyA, respectively. Finally, we compiled a user-friendly interactive application to facilitate navigation of our dataset by the scientific community (link).

**Figure 1.**
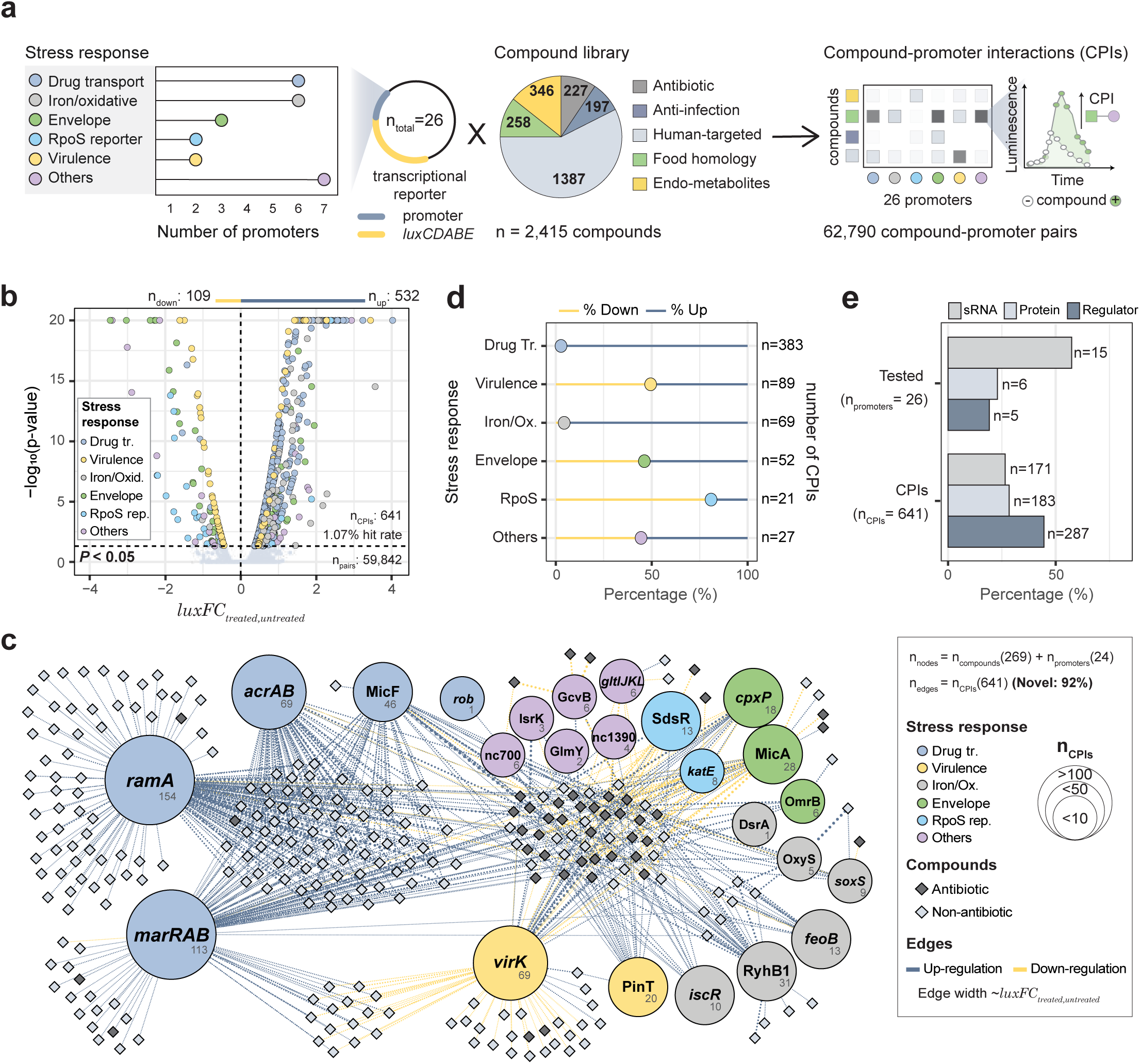
Mapping the transcriptional response of key *Salmonella* stress pathways across a large chemical space. **a)** Schematic representation of the high-throughput screening approach. The number of promoters distributed by stress response pathways and the composition of the chemical library used in this study are depicted on the left side and middle, respectively. A collection of 26 reporter strains was challenged with the 2,415 compounds, and growth and promoter signals were used to estimate CPIs. **b)** Volcano plot of all compound-promoter combinations after quality control, indicated by n_pairs_. Points represent the average of *luxFC_treated_*_/*untreated*_ and the maximum corrected p-value between two biological replicates (Methods). For ease of visualization, we have capped -log_10_(p-value) at 20. For original values see Supplementary table 4. n_CPIs_ represents the number of significant CPIs, and hit-rate reflects n_CPIs_ as a fraction of n_pairs_. The number of CPIs corresponding to up- and down-regulation of promoter activity are represented above the volcano plot, in blue and yellow, respectively. **c)** Network representation of CPIs in our dataset. Nodes represent promoters (circles) and compounds (diamonds), while edges represent the CPIs linking the promoters to the compounds. All promoters with at least one CPI are shown (24 out of 26 tested). Numbers inside the promoter nodes represent the total number of CPIs for each promoter, also reflected by node size. Percentage of novel CPIs is shown, with previously described CPIs marked in Supplementary table 5. **d)** Different profile and number of CPIs across stress responses. Number of CPIs are grouped by stress response, further differentiated between up- and down-regulation. **e)** Transcriptional regulators account for the largest fraction of CPIs. Percentage of tested promoters (top) and CPIs (bottom) distributed among the functional class of the gene (sRNA, protein or transcriptional regulator). n indicates the total number of promoters/CPIs per class.

## Results

### Mapping the transcriptional response of key *Salmonella* stress pathways across a large chemical space

We screened transcriptional responses of 26 *Salmonella* promoters following treatment with 2,415 compounds, yielding 62,790 unique compound–promoter pairs (Fig. 1a). To do so, we constructed a collection of luciferase transcriptional reporter strains of the 26 genes representative of major stress response pathways with prior evidence of transcriptional change across stress conditions^13,44^. In total, our reporter collection comprised 5 transcriptional regulators, 5 effector proteins and 16 sRNAs distributed between drug transport, virulence, iron homeostasis and oxidative stress, envelope stress, RpoS regulation, and other targets, including two sRNAs of unknown function (Fig. 1a, Supplementary table 1). The compound library covered a large chemical space, containing FDA approved drugs, including antibiotics, anti-infectives and human-targeted drugs, human endogenous metabolites and food homology chemicals (Fig. 1a, Supplementary table 2). Briefly, transcriptional change in response to each individual compound was periodically monitored by luminescence in liquid culture in biological duplicates, alongside growth (optical density, OD_600nm_, Methods). Plasmid reporters were confirmed not to decrease *Salmonella* growth (Fig. S1a). As expected, the growth of *Salmonella* was also unaffected by most compounds (Fig. S1a). Replicate correlation was overall very high (Pearson R ≥ 0.9, Fig. S1b,c), even though promoters with the lowest and highest luciferase signals were slightly noisier (Fig. S1d). Basal luminescence signal of our reporter strains (no chemical challenge) generally recapitulated previously published transcript abundance^13,44^ (Pearson R = 0.63, Fig. S1e), attesting the suitability of our assay to quantify promoter activity. Data corresponding to instances where growth inhibition by the compound was greater than 80% (mostly antibiotics) or non-concordant between the two replicates, was excluded from further analysis (n=99 compounds, Supplementary Table 3, Methods). Moreover, 10 additional compounds were excluded due to substantial interference with the reporter system, as seen by altered transcriptional activity of a promoter-less reporter (EVC, Fig. S1f, Supplementary Table 3, Methods). In total, 59,842 unique compound-promoter pairs passed our stringent quality control (Supplementary Table 4), which can be readily explored by the community through our user-friendly interactive application (link).

We then defined compound-promoter interactions (CPIs) as specific compound-mediated effects on reporter activity as previously described, with minor adjustments^38^ (Methods). Briefly, the strength of each interaction was calculated as a growth-normalized, log_2_-transformed luminescence fold-change between treatment and untreated controls, including a penalization for EVC changes to account for potential unspecific effects of each compound on the reporter system (*luxFC_treated_*_/*untreated*_, Fig. S1f, Methods). Positive and negative *luxFC_treated_*_/*untreated*_ reflect increased and decreased promoter activity, respectively (Fig. S1g). CPIs were called when *luxFC_treated_*_/*untreated*_ of treated samples fell significantly outside the distribution of untreated controls (Methods, Supplementary Table 4). We report 641 CPIs (1.07% of unique compound-promoter pairs after quality control), involving 269 compounds and 24 out of the 26 screened promoters (Fig. 1b). To our knowledge >90% of all CPIs are novel (Fig. 1c, Supplementary Table 5). The majority of the identified CPIs represent increased promoter activity, with promoters involved in drug transport and virulence responding to a greater number of compounds (Fig. 1d). For instance, ∼60% of total CPIs correspond to increased activity of a drug transport-associated promoter, mainly *ramA* or *marRAB,* encoding transcriptional regulators. On the opposite end, transcriptional activity of sRNAs is modulated by a smaller number of compounds, supporting a rather specific regulation of these elements (Fig. 1e). While most CPIs increased transcriptional activity, four promoters alone accounted for 71% of CPIs leading to downregulation, namely, the promoters of the virulence gene *virK*, the RpoE-regulated sRNA MicA, and the RpoS-regulated genes *katE* and SdsR (Fig. 1c).

A rarefaction analysis confirmed adequate sampling depth for our screen with respect to the probed chemical space. The number of promoters involved in CPIs reached saturation across the library of 2,415 compounds, indicating sufficient chemical diversity to elicit transcriptional changes across the tested promoters (Fig. S2a). Conversely, the number of compounds associated with CPIs across the 26 promoters did not saturate (Fig. S2b). This was expected as our experimental design aimed to saturate chemical space for a defined promoter set, rather than exhaustively map genome-wide responses. Furthermore, pairwise molecular structure comparisons between our library and the >12,000 small molecules in DrugBank showed that >80% of DrugBank compounds have a close structural analogue in our library (Jaccard distance between Morgan Fingerprints <0.7; Fig. S2c–d, Supplementary table 6, Methods), confirming that our library is broadly representative of the molecular diversity of the DrugBank^45^.

We then selected a subset of 45 compounds and performed a concentration-resolved validation experiment, where we re-tested each compound against each reporter strain at increasing concentrations up to 150 µM (Fig. S3a, Methods, Supplementary table 7). A Receiver Operating Characteristic (ROC) curve comparing this benchmarking set with the results of the large screen, shows a very good agreement between the two sets at the same compound concentration of 30 µM (Fig. S3b, ROC-AUC=0.90). Interestingly, ROC curve performance decayed at higher compound concentrations, which elicit stronger effects and generate additional CPIs (Fig. S3c). This points to a dose-dependent response of the promoters to the compounds, with new CPIs being uncovered when higher concentration is sampled. Altogether, this analysis shows very high reproducibility of our screen, as well as a good sampling depth of promoter activity across the tested compounds.

### Combining genome- and chemical-wide perturbation data enables the identification of chemical inducers of an out-of-screen promoter

We next benchmarked our CPI network against a publicly available transcriptomic data profiling 13 infection-relevant conditions, including macrophage environment, low pH, and cold/heat shock, among others – SalComMac^13,46^. For each dataset, we computed pairwise promoter similarity across all conditions using Pearson correlation and compared promoter-pair correlations between datasets (Fig. S4a,b; Methods). High correlation values indicate promoters that respond similarly across conditions. Strikingly, promoter pairs that correlate strongly across the diverse chemical perturbations in our dataset do not generally correlate across the 13 stress conditions, and vice versa (R = 0.11, Fig. 2a). This suggests that the two approaches – probing either a broad chemical space across a limited gene set or a broad genetic landscape across limited conditions – capture different biological insights. For instance, our dataset revealed a strong correlation between *ramAp* and *acrABp* (R = 0.85) that was weaker in the transcriptomic dataset (R = 0.31), likely because the latter did not include the conditions that induce expression of these genes (Fig. 2a and Fig. S4c).

**Figure 2.**
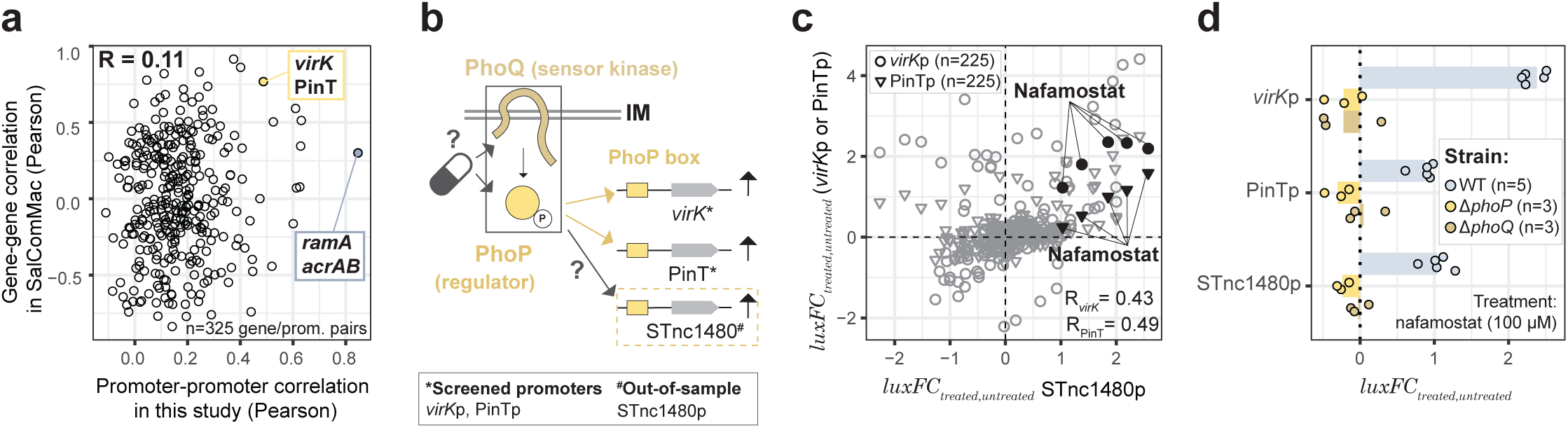
Combining genome- and chemical-wide perturbation data enables identification of chemical inducers of an out-of-screen promoter. **a)** Genome- and chemical-wide perturbations provide different biological insight. Scatter plot showing Pearson correlation coefficients between all promoter pairs from our dataset against Pearson correlation coefficients between gene expression profiles across 13 infection-relevant conditions from SalComMac ^44^. Only the genes corresponding to promoters included in our screen are represented. Pearson correlation coefficients between all promoter pairs from our dataset were calculated using *luxFC_treated_*_/*untreated*_ across 2306 compounds (passed quality control). Pearson correlation coefficients between the 26 genes from SalComMac were calculated using transcript fold-change across 13 infection-relevant conditions (explicit in Fig. S4b). R represents the Pearson correlation between the two datasets. Two examples are highlighted - *virK*-PinT as a concordant pair, and *ramA*-*acrA* as a less concordant pair, between both datasets. **b)** Schematic illustration of PhoPQ mediated regulation affecting the expression of virulence-associated promoters *virK*p, PinTp and STnc1480p. **c)** Nafamostat induces *virK*p, PinTp and STnc1480p. Scatterplot shows *luxFC_treated_*_/*untreated*_ of STnc1480p against *virK*p or PinTp across 45 compounds at 5 different concentrations (30-150 µM). All concentrations are shown individually (n=45×5=225). The 5 concentrations of nafamostat are highlighted for both promoters. Data shown corresponds to mean *luxFC_treated_*_/*untreated*_ across 4 biological replicates. R represents *luxFC_treated_*_/*untreated*_-based Pearson correlation between STnc1480p and the indicated promoter. **d)** Nafamostat induces *virK*p, PinTp and STnc1480p in a PhoPQ-dependent manner. n represents the number of biological replicates, independently shown as dots. Bars represent the average across the indicated biological replicates. *luxFC_treated_*_/*untreated*_ reflects the fold-change of promoter activity in response to treatment, and it was calculated as described in Methods (**a, b** and **d**).

Despite their apparent orthogonality, we hypothesized that the integration of the two datasets could yield biological insights not accessible from either alone. To test this, we focused on the virulence-associated genes *virK* and PinT, which show moderate-to-strong correlation in both datasets (Fig. S4d). The transcriptomic data shows that additional genes correlate with *virK* and PinT, including the sRNA STnc1480, whose promoter was not included in our dataset (Fig. S4e). Consistent with their correlated transcriptional activity, all three genes are known to be co-regulated by the PhoPQ two-component system^46–48^. Our screen identified over 30 compounds – primarily, but not exclusively, antibiotics – that increase activity of either *virK* or PinT (Fig. 1c, Supplementary Table 5). Therefore, we hypothesise that the same compounds may also increase promoter activity of STnc1480, possibly via PhoPQ regulation in all three cases (Fig. 2b). To test this hypothesis, we measured STnc1480p promoter activity across the 45 compounds used for the screen validation above, and compared its activity profile with that of *virK*p and PinTp (Methods, Supplementary Table 7). STnc1480 promoter activity moderately correlated with *virK*p and PinTp (Fig. 2c), suggesting a substantial fraction of compounds that co-regulate the three genes. One such compound with no previous association with the PhoPQ two component system is the human drug nafamostat (Fig. 2c), a prescription-bound serine-protease inhibitor used as an anticoagulant^49^. We then established that nafamostat increases the activity of the three promoters in a PhoPQ-dependent manner, since deletion of either *phoP* or *phoQ* abolished this effect (Fig. 2d). Interestingly, it has been shown that these three promoters show tight co-regulation by 19 regulators, beyond PhoP^46^ (Fig. S4f), and thus it remains to be investigated whether nafamostat exerts its effects exclusively via PhoP, or also via other regulatory programs. Altogether, besides exposing a novel chemical inducer of PhoPQ regulated genes, this example showcases how combining our dataset with available transcriptomic data can enable us to expand our CPI network to include promoters originally absent from our screen.

### Antibiotics and non-antibiotic compounds elicit distinct regulatory stress responses in *Salmonella*

Antibiotics extensively modulate gene expression in bacteria^31,38,50^. Consistent with this, they are enriched among CPIs in our dataset, even though human-targeted drugs account for nearly half of all detected interactions (Fig. S5 a,b). Additionally, antibiotics participate in CPIs with a greater number of promoters than other anti-infective drugs, food homology, human-targeted compounds or endogenous metabolites (non-antibiotics, Fig. 3a). Concomitantly, antibiotics also inflict a larger growth defect than other compound classes in our dataset (Fig. 3b). Altogether, this suggests that CPIs involving antibiotics may be non-specific, and possibly driven by compound impact on growth, rather than direct regulation of a given promoter. In contrast, human-targeted drugs differ fundamentally from antibiotics, as they mostly did not affect growth in our dataset, and modulate the expression of fewer promoters (Figs. 3a,b), consistent with a more specific and potentially direct regulatory scope.

**Figure 3.**
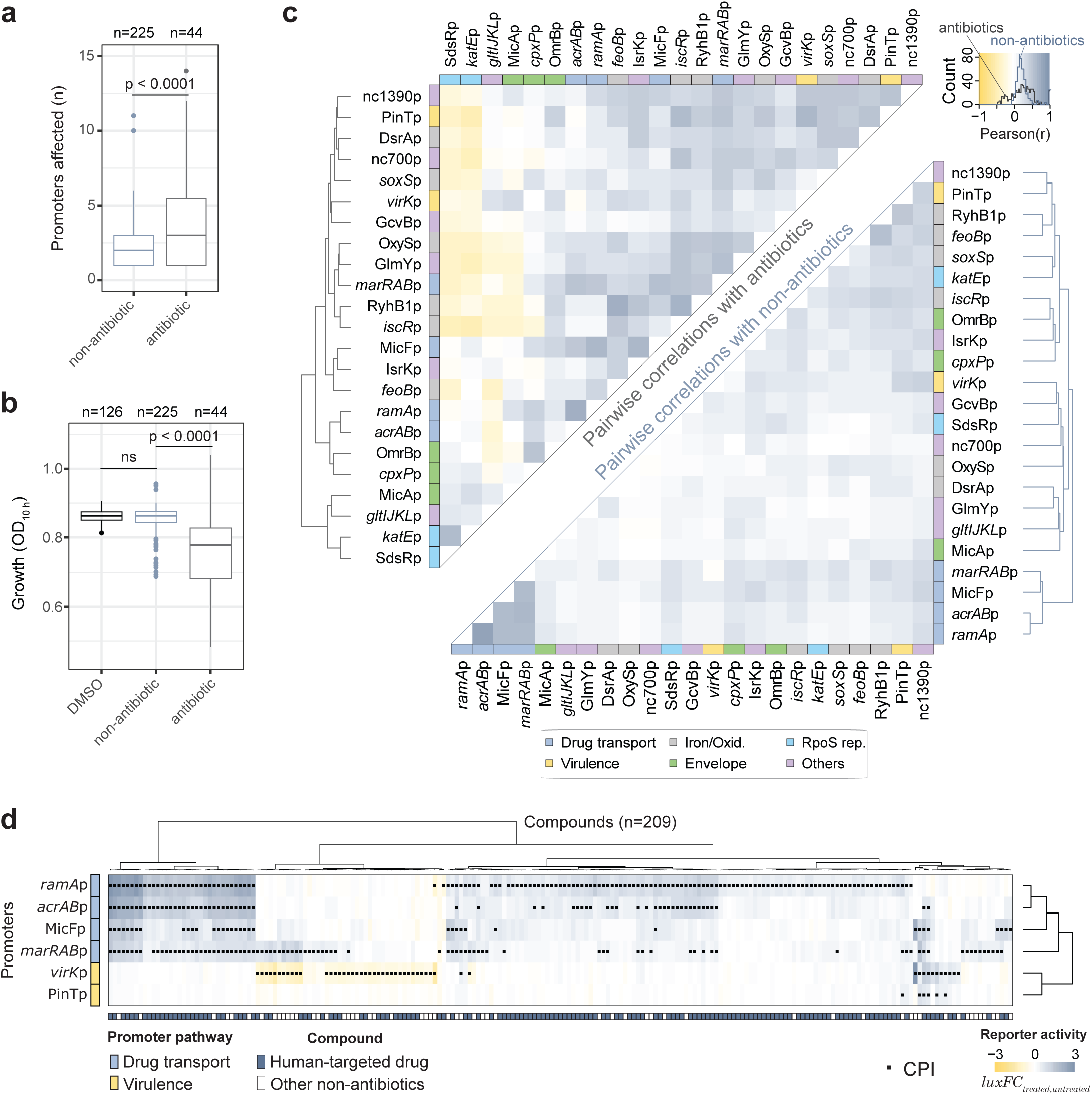
Antibiotics and non-antibiotic compounds elicit distinct regulatory stress responses in *Salmonella*. **a)** Antibiotics affect a higher number of promoters. Boxplots show the number of promoters involved in CPIs with either antibiotics or non-antibiotics. n reflects the number of antibiotics or non-antibiotic compounds involved in CPIs in our dataset. The p-value of a double-sided Wilcoxon test between the two distributions is shown. Boxplots indicate 25th, 50th and 75th percentiles, and whiskers extend up to 1.5 times the interquartile range (IQR) from the 25th and 75th percentiles (applies to **a** and **b**). **b)** Non-antibiotics generally do not reduce *Salmonella* growth. Boxplots show Salmonella growth for untreated wells (DMSO), non-antibiotics and antibiotics. n reflects the number of antibiotics or non-antibiotic compounds involved in CPIs in our dataset, and all untreated control wells. Growth with each compound was estimated as the mean growth of all reporter strains (n=26 + EVC), where the value of each strain was first averaged from the two biological replicates. **c)** Antibiotics and non-antibiotics elicit distinct regulatory responses. Correlation matrices between promoter pairs across antibiotic (n=130, top) and non-antibiotic (n=2176, bottom) compounds. Color intensity reflects *luxFC_treated_*_/*untreated*_-based Pearson correlation coefficients. The dendrograms show hierarchical clustering based on Euclidean distances with complete linkage. **d)** Drug transport promoters and the promoter of *virK*, a virulence protein, account for most CPIs involving non-antibiotics. A matrix showing *luxFC_treated_*_/*untreated*_ of drug transport and virulence-associated promoters across 209 non-antibiotics is shown. All non-antibiotic compounds involved in at least one CPI with the depicted promoters are included. Significant CPIs are indicated as dots. The dendrograms show hierarchical clustering based on Spearman correlation and complete linkage for the promoters, and Euclidean distances with Ward’s criterion for compounds.

We next assessed similarity in promoter responses to antibiotic and non-antibiotic compounds by computing pairwise Pearson correlations, generating separate promoter–promoter correlation matrices for each group. (Fig. 3c, Methods). Both correlation matrices were subjected to hierarchical clustering, allowing promoters with similar compound-dependent response patterns to group together. This analysis revealed that clustering of promoters differed between antibiotics and non-antibiotics, with generally stronger correlations observed under antibiotic treatment (Fig. 3c). For instance, canonical RpoS-dependent stress responses cluster together under antibiotic treatment (Fig. 3c), and are anti-correlated with most other stress reporters, recapitulating previous findings^51,52^ (Fig. S5c). By contrast, drug transport genes cluster together under non-antibiotics, but not under antibiotic treatment, suggesting different modes of regulation of these promoters by these two sets of compounds (Fig. 3c). In fact, we observed that non-antibiotic compounds mostly trigger the response of drug transport- and virulence-associated genes, in particular of the outer membrane virulence determinant *virK* (Fig. 3d and S5d). This distinctive and poorly understood transcriptional response to non-antibiotics prompted us to further investigate its molecular basis.

### The activity of the virulence regulator SlyA is modulated by non-antibiotic compounds

VirK is a virulence-associated protein that also contributes to resistance against cationic antimicrobial peptides in several Gram-negative pathogens, including *Salmonella, Shigella*, *E. coli* and *Klebsiella pneumoniae*^47,53,54^. In our screen, we identified 38 non-antibiotic compounds that decrease activity of the *virK*p promoter (Supplementary Table 3, Fig. 4a), including several human metabolites, hydrocinnamic acid derivatives typically found in food, and non-steroidal anti-inflammatory drugs (NSAIDs). Notably, these compounds did not affect expression of the other PhoPQ-regulated promoter PinTp, suggesting that interference with PhoPQ signalling is unlikely to explain the observed downregulation of *virK* (Fig. 3d). To further investigate this effect, we focused on dantrolene, a calcium channel antagonist commonly used as a muscle relaxant ^55^, which strongly reduced *virK*p activity. Dantrolene completely overruled the canonical PhoPQ response of *virK*p: *virK*p activity remained suppressed by dantrolene regardless of Mg²⁺ concentration in the medium (Fig. 4b). Moreover, the inhibitory effect of dantrolene persisted in a Δ*phoP* mutant, conclusively ruling out PhoPQ-mediated regulation (Fig. 4b).

**Figure 4.**
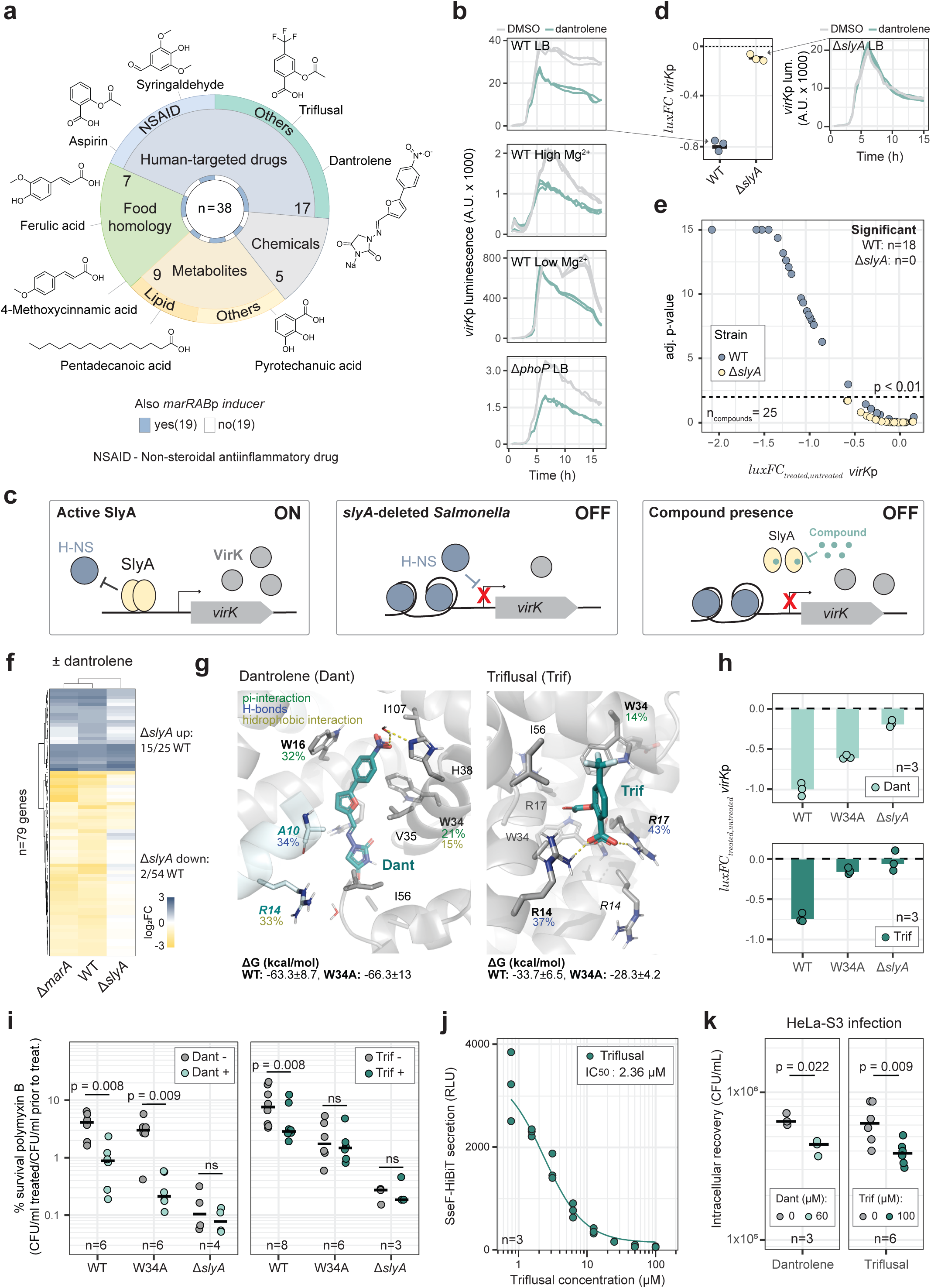
The activity of the virulence regulator SlyA is modulated by non-antibiotic compounds. **a)** Overview of the non-antibiotics that down-regulate *virK*p. **b)** Dantrolene down-regulates *virK*p independently of PhoPQ. *virK*p luminescence-over-time curves of untreated (DMSO) and dantrolene-treated *Salmonella* wild type (WT) or Δ*phoP* across different Mg^2+^ concentrations are shown. High Mg^2+^ and Low Mg^2+^ indicate LB medium supplemented with 10 mM of MgCl_2_ and N-minimal media supplemented with 10 µM of MgCl_2_, respectively. Three biological replicates are shown. **c)** Schematic representation of the proposed model of compound-mediated *virK* regulation via SlyA. In basal conditions (left), SlyA facilitates *virK* expression by binding to its promoter region, thereby counteracting silencing by H-NS ^37^. In the absence of SlyA (middle), H-NS binds to the *virK* promoter region, thereby reducing its expression. We hypothesize that dantrolene, and possibly other compounds in our dataset, cause SlyA to release the DNA, thereby allowing *virK* silencing by H-NS (right). **d)** Dantrolene mediated downregulation of *virK*p is *slyA* dependent. Left: Dotplot showing the quantification of *luxFC_treated_*_/*untreated*_ *virK*p based on the luminescence-over-time curves of untreated (DMSO) and dantrolene-treated *Salmonella* wild type (WT) and Δ*slyA*. Right: *virK*p luminescence-over-time curve of untreated (DMSO) and dantrolene-treated *Salmonella* Δ*slyA.* Data from three biological replicates is shown. **e)** Non-antibiotics downregulate *virK*p via SlyA. Volcano plot showing *virK*p *luxFC_treated_*_/*untreated*_ of wild type and Δ*slyA Salmonella* across 25 non-antibiotics that decreased *virK*p activity in our dataset (subset from 38 shown in **a**). All experiments were done at 100 µM, in three biological replicates. Mean *luxFC_treated_*_/*untreated*_ across biological replicates is shown. The p-value of a double-sided Z-test comparing three biological treated samples with 48 untreated controls after correction for multiple testing (Benjamini-Hochberg) is shown. For ease of visualization, we have capped -log_10_(pvalue.adj) at 15. **f)** SlyA mediates most transcriptional adjustments caused by dantrolene in *Salmonella*. Heatmap shows the transcriptional log_2_FC of the 79 differentially expressed genes in wild type *Salmonella* upon dantrolene treatment. Data for wild type, Δ*marA* and Δ*slyA* is shown, and it reflects changes across 3 biological replicates. Dendrograms reflect hierarchical clustering based on Euclidean distance and complete linkage. **g)** Triflusal and dantrolene bind to SlyA *in silico*. Representative binding mode for triflusal (left) and dantrolene (right) on different pockets derived from molecular dynamic (MD) simulations. MD trajectories of 5×300-500 ns were clustered by the ligand’s heavy atoms position variation to retrieve the representative binding mode. Interaction frequencies along the simulations are depicted as numbers near the amino acid labels (green – pi-mediated interactions, blue – polar contacts and yellow – hydrophobic interactions). Predicted binding energy values for triflusal and dantrolene calculated along the simulation time for SlyA and SlyA-W34A are depicted below. **h)** W34A mutation in SlyA hinders triflusal-but not dantrolene-mediated inhibition of *virK*p. Barplots showing mean *luxFC_treated_*_/*untreated*_ *virK*p for wild type, *slyA*-W34A and Δ*slyA* upon treatment with dantrolene or triflusal (100 µM). Data from three biological replicates is shown. **i)** SlyA-inhibitors increase sensitivity to polymyxin B. Dotblots showing survival of *Salmonella* wild type, *slyA*-W34A and Δ*slyA* to polymyxin B treatment, with and without dantrolene (left) or triflusal (right). Horizontal lines represent the mean CFU/ml across n biological replicates. p-values from double sided paired t-tests are shown directly above the compared samples. **j)** Triflusal reduces activity of the type-III secretion system encoded in SPI-2 (T3SS-2). The dose-response curve of SseF-HiBiT secretion by T3SS-2 with increasing triflusal concentrations is shown. Secretion activity was measured as previously established ^62^. n indicates the number of biological replicates. **k)** SlyA-inhibitors impair *Salmonella* intracellular survival in HeLa-S3 cells. Dotblots showing survival of *Salmonella* wild type upon recovery from infected HeLa-S3 cells with and without dantrolene (left) or triflusal (right). Compound treatment was performed after HeLa-S3 cells invasion in a gentamicin protection assay. Time of infection = 8 hours, Multiplicity of Infection (MOI) = 20. Horizontal lines represent the mean CFU/ml across n biological replicates. p-values from double sided paired t-tests are shown directly above the compared samples. Dantrolene and triflusal concentrations are indicated in the legend.

Interestingly, nearly half of the compounds that decreased *virK*p activity also increased activity of the drug transport-related promoter *marRAB*p, which is primarily repressed by the MarR-family repressor MarR (Fig. 3d, Fig. 4a). While MarR has not been linked to *virK* regulation, the related protein SlyA, also a MarR-family regulator, is a well-established regulator of virulence genes in *Salmonella*, including *virK*^56–58^. Thus, we hypothesized that SlyA mediates the downregulation of *virK* by non-antibiotic compounds in our dataset (Fig. 4c). Consistent with this hypothesis, dantrolene-mediated inhibition of *virK*p was abolished in a Δ*slyA* mutant (Fig. 4d). Expanding this analysis to 25 additional non-antibiotic compounds that decreased *virK*p activity in our dataset confirmed significant downregulation for 70% of them, all in a *slyA*-dependent manner (Fig. 4e, Fig. S6a, Supplementary table 8). These results strongly support SlyA as the primary regulator of non-antibiotics-induced repression of *virK* in our dataset.

Given the broader regulatory role of SlyA, we next examined the global transcriptional response to dantrolene, a representative of non-antibiotics that decrease *virK*p promoter activity. Dantrolene induced widespread transcriptional changes, with 79 differentially expressed genes, including 54 downregulation events (Fig. S6b, Supplementary Table 9). Among these was the SlyA-regulated gene *pagC*^57^. Although *virK* fell just below the stringent statistical cutoff in this experiment, we confirmed its downregulation by RT-qPCR (Fig. S6c). Notably, dantrolene also increased expression of the *marRAB* and *emrAB* operons, both normally repressed by MarR-family regulators under non-inducing conditions (MarR and EmrR, respectively, Fig. S6b). These results suggest that dantrolene modulates the activity of multiple MarR-family regulators, including SlyA, MarR, and EmrR. To disentangle the contribution of SlyA from that of MarR, we re-evaluated the effect of the drug in the Δ*slyA* and Δ*marA* mutants (the latter hinders genome-wide regulation by MarR). Deletion of *slyA* markedly attenuated the transcriptional response observed in wild-type cells, with more than 90% of the downregulated genes losing their response (Fig. 4f). In contrast, *marA* deletion of had only a minor effect on the transcriptional response to dantrolene, indicating that SlyA is the primary regulator mediating this response (Fig. 4f). Interestingly, deletion of *slyA* abolished dantrolene-induced overexpression of *marRAB* in this dataset (Fig. S6d), questioning the function of the MarR repressor in regulating the *marRAB* operon itself in these conditions. Indeed, deletion of *slyA* had a more substantial impact on *marRAB*p promoter activity with and without dantrolene than deleting *marR* itself (Figs S6e,f). Altogether, these results challenge the well-established role of a single-headed MarR-dependent regulation in response to chemical signals^59^, and rather highlight a prominent role of SlyA, and an intricate cross-talk between MarR-family regulators.

### Computational structural analysis suggests compound-mediated regulation by direct SlyA binding

Our results so far show that dantrolene, and presumably other non-antibiotics that decreased *virK*p promoter activity, inhibit SlyA function, and therefore we now refer to them as SlyA-inhibitors. We then hypothesised that inhibition could be caused to direct binding to SlyA. Indeed, some of the identified SlyA-inhibitors have similar chemical structures to established SlyA binders^35,60^, e.g. triflusal (a platelet aggregation inhibitor, Fig. 4a). However, other SlyA-inhibitors such as dantrolene have substantially different chemical structures, which have not yet been shown to bind MarR-regulators (Fig. 4a). Hence, we used molecular dynamic simulations to assess the likelihood of the SlyA-inhibitors, namely the structurally distinct dantrolene and triflusal, to bind SlyA. The simulations predicted that both compounds associate with SlyA with binding energies comparable to those of the positive control, salicylate (Supplementary information), supporting a similar binding propensity (Fig. 4g).

In agreement with previous analysis^61^, we predicted that the tryptophan at position 34 (W34), as well as the arginine residues at positions 14 and 17 (R14 and R17) contribute to SlyA ligand-binding. Thus, we generated three SlyA variants, each harbouring one of these point mutations (SlyA-W34A, SlyA-R14A, SlyA-R17A), and tested their impact on *virK*p promoter activity, with and without SlyA-inhibitors. In agreement with previous observations^61^, we found that in the absence of the inhibitors, SlyA-W34A regulatory activity was only slightly reduced compared to SlyA, while mutations R14A and R17A completely abolished SlyA function, and were therefore excluded from further analyses (Fig. S6g). In the presence of the inhibitors, we found that triflusal-, but not dantrolene-, mediated inhibition of *virK*p was significantly attenuated in the SlyA-W34A mutant (Fig. 4h), reflecting the structural differences and potentially distinct binding properties of the two compounds (Fig. 4g and h). Further molecular docking simulations confirmed that SlyA-W34A binding to triflusal becomes slightly less energetically favourable, while the energy state of dantrolene bound SlyA-W34A is unaffected (Fig. 4g).

### SlyA-inhibitors increase sensitivity to polymyxin B and decrease *Salmonella* virulence in cell culture

Deletion of *slyA*, as well as of SlyA-regulated genes, such as *virK* and *pagC*, have been previously shown to increase *Salmonella* sensitivity to the membrane targeting antibiotic polymyxin B^47,58^. We therefore hypothesised that SlyA-inhibitors could have a similar effect. Under conditions in which VirK regulation by SlyA regulation is most impactful (low Mg^2+^), dantrolene and triflusal enhanced bactericidal activity of polymyxin B (Fig. 4i). This phenotype was abolished in the *slyA*-W34A treated with triflusal, further supporting that ligand-binding by SlyA is likely to occur and drive the phenotype (Fig. 4i). By contrast, dantrolene-mediated enhanced killing by polymyxin B was not altered in the SlyA-W34A mutant, further supporting that the residue W34 is not decisive for ligand-binding by SlyA in the presence of dantrolene. Importantly, deletion of *slyA* completely abolished the ability of both compounds to enhance the bactericidal activity of polymyxin B (Fig. 4i).

As SlyA is a central regulator of *Salmonella* pathogenicity, namely of type III secretion system (T3SS-2) encoded in *Salmonella* Pathogenicity Island 2 (SPI-2)^56^, we assessed whether SlyA-inhibitors decrease secretion of virulence factors by this system. Using a previously described secretion reporter^62^, we found that triflusal strongly reduces T3SS-2 secretion activity in a concentration dependent manner (Fig. 4j). The nature of the assay, in particular acidic conditions, precluded us to assess the effect of dantrolene on SPI-2 secretion, due to its chemical instability in these conditions^55^ (Fig. S6h). Subsequently, we assessed if the same SlyA inhibitors impaired *Salmonella* virulence *in vitro*, and observed that both dantrolene and triflusal significantly reduced intracellular *Salmonella* recovered upon infection of HeLa cells (Fig. 4k). Importantly, none of these compounds displayed cytotoxic effects (Methods, Fig. S6i). Altogether, our results expose the virulence regulator SlyA as a major regulator of chemical stress in *Salmonella*, with potential for anti-infective applications.

### Human-targeted drugs increase intrinsic antibiotic resistance via RamR-RamA in *Salmonella*

Having identified a major regulator mediating the response of virulence-associated promoters to non-antibiotics, we sought to further characterize CPIs involving non-antibiotics and drug transport-associated promoters. We identified 206 compounds in CPIs with drug transport-associated promoters (Supplementary table 5), 180 of which are non-antibiotics including human-targeted drugs of broad applications, as well as antifungal and antimalaria compounds (Fig. 3d, Fig. 5a). Notably, 48% of all non-antibiotics that increased activity of at least one drug transport promoter, increased activity of at least another (Fig. S7a), upholding the regulatory crosstalk between the promoters in these conditions. The unprecedented number of human-relevant molecules found to coordinate the transcription of promoters involved in bacterial drug resistance prompted us to investigate the molecular underpinnings of this phenomenon. As well-established, environmentally responsive multidrug-resistance transcription activators in *Enterobacteriaceae*, both MarA and RamA are well positioned to coordinate such regulatory networks^31,38^. Therefore, we selected 16 non-antibiotics that triggered expression of at least one drug transport-associated promoter (Fig. S7b), and tested promoter activation in Δ*ramA* and Δ*marA* mutants. Remarkably, none of the deletion mutants showed impaired growth with the selected compounds (Fig. S7c). We found that *ramA*, but not *marA*, is required for activation of the drug transport-associated promoters *acrAB*p, MicFp, *marRAB*p, and *ramA*p by 15 of the 16 non-antibiotic compounds tested (Fig. 5b). The exception was dantrolene, which acts through SlyA and was included as a negative control. (Fig. 5b). Furthermore, we observed high correlation between promoter activities of all drug transport-associated promoters with *ramA*p across the 139 non-antibiotics involved in CPIs with *ramA*p (Fig. S7d), establishing RamA as a central regulator of intrinsic drug resistance promoters mediated by non-antibiotics in *Salmonella*.

**Figure 5.**
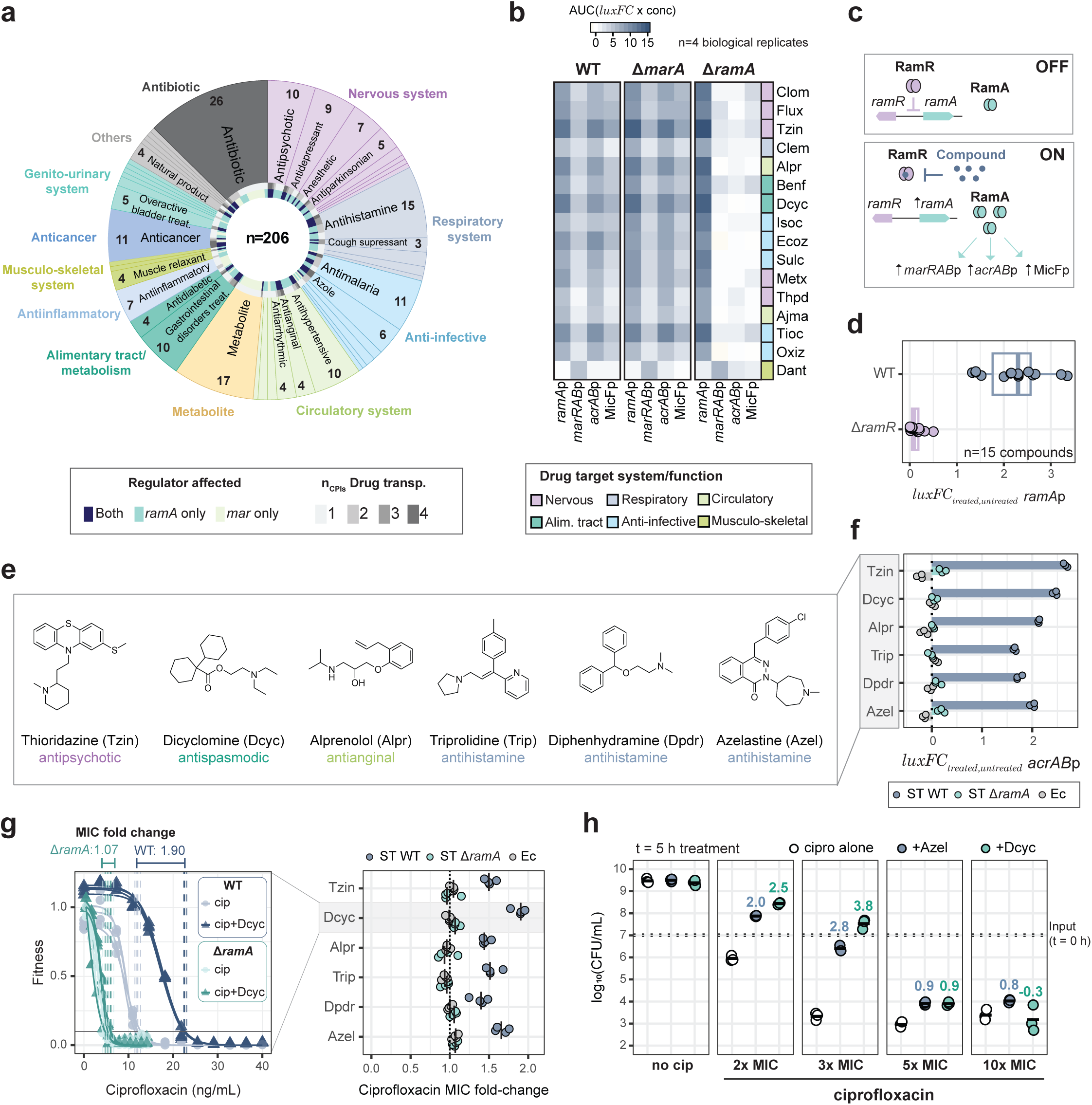
Widespread induction of antibiotic resistance promoters by human-targeted drugs is mediated by RamA in *Salmonella.* **a)** Overview of all compound classes involved in CPIs with drug transport-related promoters. Aside from drug function, the number of drug transport promoters (shades of gray) and which of the regulator-encoding promoters (*ramA*p – teal, *marRAB*p – light green and both – dark blue) are affected by each compound are also shown. **b)** Non-antibiotic-mediated induction of drug transport promoters is *ramA*-dependent. Heatmaps showing promoter activity of the four drug transport promoters in our dataset over a concentration range of 16 compounds selected from **a**, for *Salmonella* wild type (left), *ΔmarA* (middle) and *ΔramA* (right). Promoter activity is shown as area under the curve of *luxFC_treated_*_/*untreated*_ over increasing compound concentrations (30 to 150 µM). Color intensity reflects promoter activity averaged across n=4 biological replicates. The colors on the right side reflect the drug targeted process as in **a**. **c)** Schematic illustration of compound-mediated *ramA* regulation via RamR. In basal conditions, RamR binds to the ramA promoter region, thereby repressing its expression (top). Upon compound binding, RamR is released from the DNA and *ramA* expression is enabled, leading to downstream induction of *marRAB*p, *acrAB*p and MicFp (bottom). **d)** *ramR* mediates increased *ramA* expression by non-antibiotics. Boxplots showing *ramAp* promoter activity upon treatment with 15 compounds from **b** (except dantrolene) in *Salmonella* wild type *ΔramR*. The mean promoter activity of each individual compound across three biological replicates is shown as overlaid dots. All compounds were used at 100 µM. Boxplots indicate 25th, 50th and 75th percentiles, and whiskers extend up to 1.5 times the interquartile range (IQR) from the 25th and 75th percentiles. Luminescence curves of *ramA*p expression with the 15 compounds are shown in Fig. S7f. **e)** *ramA* inducers have varied functions and chemical structures. Chemical structure of 6 exemplary *ramA* inducing compounds that present different therapeutic applications, broad usage and high induction of the promoter of the AcrAB efflux pump. **f)** *ramA* inducers increase *acrAB* promoter activity in a *ramA* dependent manner. Barplots showing *acrAB*p promoter activity upon treatment with the 6 ramA inducers from **e** for *Salmonella* (ST) wild type *ΔramA*, as well as for *E. coli* (Ec), that naturally lacks *ramA* and *ramR*. Bars represent the average of three biological replicates. **g)** *ramA* inducers increase *Salmonella* intrinsic resistance to antibiotics in a *ramA*-dependent manner. Left: Ciprofloxacin minimum inhibitory concentration (MIC) curves for *Salmonella* wild type (blue/gray) and *ΔramA* (light/dark green) with and without 100 µM of the *ramA* inducer dicyclomine. Vertical dashed lines represent MIC of four individual biological replicates. Horizontal bars and numbers above the plots correspond to the MIC fold change between untreated and dicyclomine treated cultures. Right: Dotblots showing ciprofloxacin MIC fold change caused by the 6 ramA inducers from **f** for *Salmonella* (ST) wild type *ΔramA*, as well as for *E. coli* (Ec). Four biological replicates are shown. **h)** *ramA* inducers hinder killing by ciprofloxacin. Dotblots showing survival of *Salmonella* wild type to increasing concentrations of ciprofloxacin, with and without the *ramA* inducers dicyclomine and azelastine. Short horizontal full lines show the mean CFU/ml across three biological replicates. Horizontal dashed lines indicate the number of bacteria before treatment (input, t = 0). Bold numbers represent the average log_10_(difference) between ciprofloxacin treatment with and without *ramA* inducers.

Compound-mediated induction of *ramA* expression has been shown to rely on ligand binding to the TetR-family repressor RamR^26^ (Fig. 5c). Only a few compounds have been described to bind RamR and de-repress *ramA* expression^32,33^. Nonetheless, we hypothesised that the compounds that induce the *ramA* promoter in our dataset could do so via RamR de-repression. Indeed, we confirmed that *ramR* is strictly necessary for compound-mediated induction of *ramA* expression by the 15 human-targeted drugs selected above, strongly suggesting that these compounds could in fact act via RamR (Figs. 5d, S7e,f).

Next, we questioned whether the non-antibiotics involved in CPIs with drug-resistance promoters would increase *Salmonella* resistance to antibiotics. Thus, we selected six drugs among the non-antibiotic compounds in our dataset, namely three anti-histamines, an antipsychotic, an antianginal and antispasmodic, based on different therapeutic applications, broad usage and high induction of the promoter of the AcrAB-TolC efflux pump (Fig. 5e). We confirmed that the latter effect was dependent on the presence of *ramA*, as increased *acrAB*p activity was not observed in a Δ*ramA Salmonella* mutant and in *E. coli*, which naturally lacks *ramA* (Fig. 5f). We then assessed whether the six selected drugs influenced resistance to two different antibiotics that are well documented substrates of the AcrAB-TolC efflux pump - ciprofloxacin and chloramphenicol^9,63^. We measured minimum inhibitory concentration (MIC, minimum concentration required to fully inhibit microbial growth, Methods) of each antibiotic alone and in combination with each drug, and observed that all six drugs significantly increased the MIC of both antibiotics up to 2-fold, in a *ramA*-dependent manner (Figs. 5g and S7g). In addition, we also found that the compounds that strongly antagonised ciprofloxacin activity, dicyclomine and azelastine, also protected *Salmonella* from being killed by this antibiotic at concentrations moderately above MIC (Fig. 5h). Altogether, these results establish RamA as a broad responder to non-antibiotic drugs in *Salmonella* and showcase that RamA-mediated increased expression of drug transport genes by human-targeted drugs increases intrinsic resistance to antibiotics.

### Machine learning approach enables large-scale, out-of-sample prediction of human-targeted drugs inducing *ramA* expression in *Salmonella*

While previous studies have reported *ramA* induction and RamR binding by a number of structurally diverse molecules^30–33^, some of which we recapitulate here, our dataset expands this landscape by identifying over 130 human-targeted drugs that induce *ramA*, most of which not previously reported (Fig. 1c, Supplementary table 5). This shows that the effect of human-targeted drugs on the expression of intrinsic antibiotic resistance genes in *Salmonella* goes far beyond what was previously acknowledged, and urged us to characterise the underlying chemical landscape of *ramA* inducers. We used our unique dataset to train a machine learning model that predicts compound ability to induce *ramA* expression based on its molecular structure (Fig. 6a). Briefly, compounds were split into training, validation, and test sets based on molecular scaffolds, and a LASSO logistic regression model was trained using Morgan fingerprints as molecular features (Methods, Supplementary table 10). The model showed overall good predictive performance, with areas under the ROC and precision-recall curves of 0.77 and 0.39, respectively (Fig. S8a,b). In particular the model exhibited high precision at identifying compounds that induce *ramA* expression, as shown by its low false-positive rate (Fig. S8c). This comes at the expense of a reduced sensitivity, as the model misses up to 50% of true inducers found in our screen (considering all data - Fig. S8c). Nonetheless, the particularly low false-positive rate gave us confidence to further explore the results, namely the molecular features of *ramA* inducers.

**Figure 6.**
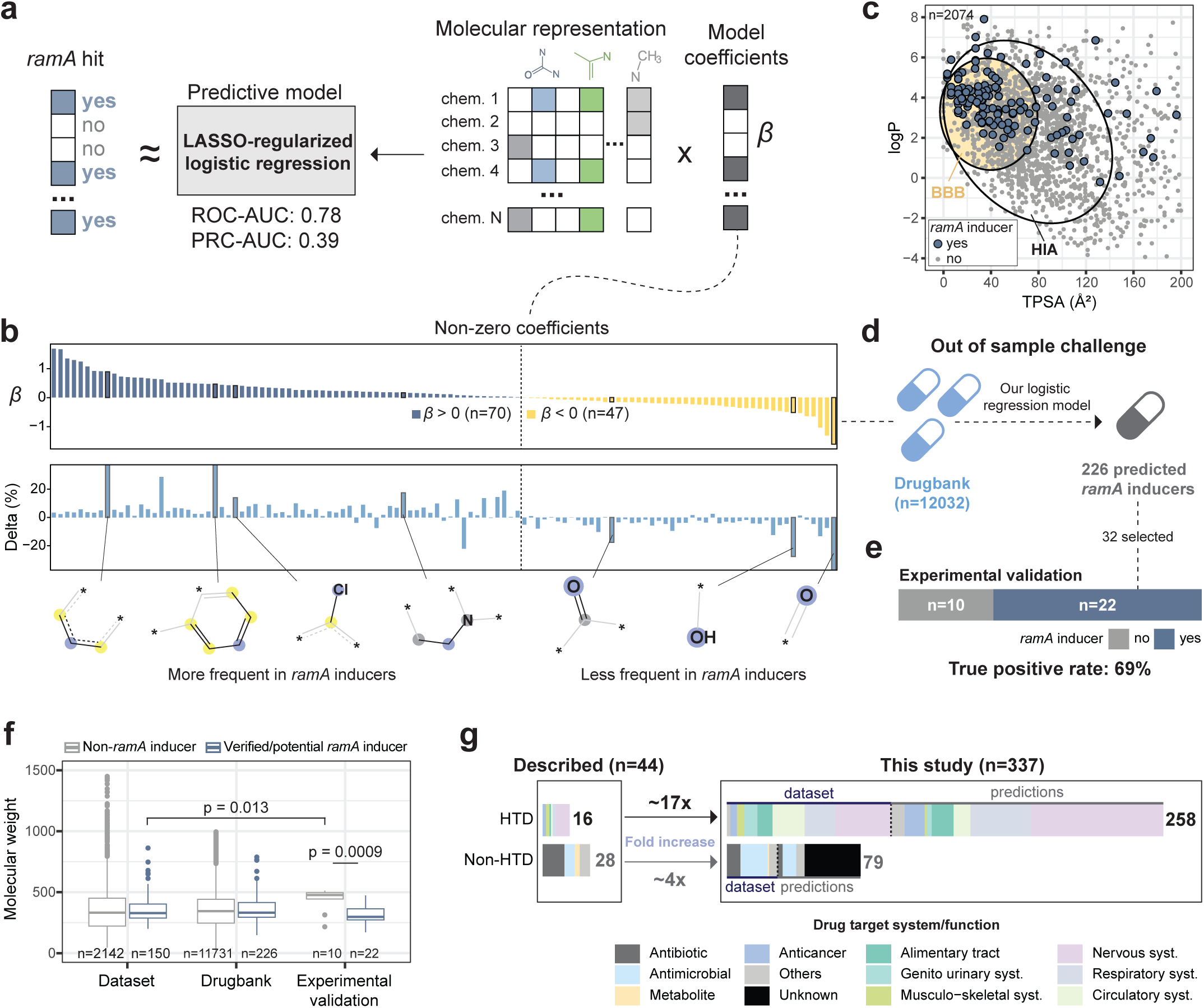
Machine learning approach enables large-scale, out-of-sample prediction of human-targeted drugs that increase *ramA* expression in *Salmonella.* **a)** Scheme illustrating the *ramA* induction predictive model. The >2000 compounds from our dataset were classified as *ramA* inducers and non-inducers. Morgan fingerprints were used for chemical representation of the compounds to train a logistic regression model for predicting *ramA* induction. The contribution of each fingerprint to prediction is reflected by model coefficients *β*. LASSO penalization was used to reduce overfitting and eliminate less informative features from the model. **b)** Molecular features that distinguish *ramA* inducers. Top: bars show non-zero model coefficients *β* of 114 features. Positive (blue) and negative (yellow) *β* reflect contribution of each feature to prediction of *ramA* inducers. n represents the number of non-zero features. Bottom: bars show the difference between the percentage of each feature among *ramA* inducers and non-inducers (Delta %). Illustration of exemplary chemical features over-(left) or underrepresented (right) in *ramA* inducers. Atom colors refer to the description provided at the RDKit documentation (blue: central atom, yellow: aromatic atom, gray: aliphatic ring atom). c) *ramA* inducers present reduced polarity and increased lipophilicity. BOILED egg plot of the compounds present in our dataset, represented by a scatterplot of logP (partition coefficient) and TPSA (topological polar surface area) for each compound. The highlighted areas represent typical logP and TPSA ranges of compounds absorbed in the human intestine (human intestine absorption, HIA, white) and compounds that cross the blood-brain barrier (BBB, yellow). **d)** Out-of-sample challenge of our model reveals additional candidate ramA inducers. The *ramA* inducer predictive model was applied to the Drugbank set of 12,032 compounds to identify additional candidate *ramA* inducers. 226 additional candidate *ramA* inducers, absent from our initial dataset, were identified. **e)** Experimental validation of out-of-sample challenge reinforces good predictive performance. 32 compounds out of newly predicted 226 *ramA* inducers were selected and tested for *ramA* promoter activity at 100 µM (Fig. S9c). Bars show the number of true *ramA* inducers and not inducers among the 32 selected. True positive rate corresponds to the percentage of true inducers among the 32 compounds tested. f) *ramA* inducers occupy a constrained range of molecular weight. Boxplots show the molecular weight of all compounds in our dataset, DrugBank, or selected for experimental validation of the out-of-sample challenge, split between non-*ramA* inducers and *ramA* inducers/candidates. Boxplots indicate 25th, 50th and 75th percentiles, and whiskers extend up to 1.5 times the interquartile range (IQR) from the 25th and 75th percentiles. p-value of double sided Wilcoxon rank-sum tests are shown. **g)** Expansion of *ramA* inducers among human-targeted drugs. Bars show the total number of *ramA* inducers reported in literature (n=44), and new compounds uncovered (dataset) or predicted (predictions) in this study (n=337), split between human-targeted drugs (HTD) and non-HTD. Only non-redundant compounds were accounted for; for instance, different salts of the same compounds were accounted for only once. Compounds previously described are exclusively accounted for as “described”, and absent from “this study”, even if we were able to re-discover them (Supplementary table 12).

Leveraging the interpretability enabled by Morgan Fingerprints, we investigated the most informative chemical features that predict *ramA* induction in our model. Out of the 2048 original features, 117 were found to contribute to *ramA* induction (Fig. 6b, Fig. S8d, Supplementary table 11, Methods). Moieties containing aromatic rings, tertiary amines and aromatic ring-substituted heteroatoms such as chlorine are enriched among the *ramA* inducers (Fig. 6b). This aligns well with molecules previously identified to induce *ramA* expression by directly binding to RamR^32^. Importantly, our model also revealed a set of common chemical features under-represented in *ramA* inducers, such as carbonyl and hydroxy groups (Fig. 6b). Altogether, these features suggest that *ramA* inducers could have certain physicochemical properties, namely high lipophilicity. Since high lipophilicity is strongly associated with human-targeted drugs, especially those crossing the blood-brain barrier, we asked whether *ramA* inducers feature common physicochemical properties with these drugs. We computed the lipophilicity and topological polar surface area (logP and TPSA, Methods) of the 2,415 compounds tested in this study, and confirmed that most *ramA* inducers indeed exhibit physicochemical properties typical of drugs crossing the blood-brain barrier^64^ (Fig. 6c). Notably, many other drugs display similar properties, but do not induce *ramA*, highlighting that physicochemical properties alone may not be sufficient to predict *ramA* induction. Indeed, when we trained an alternative model accounting only the physicochemical properties of each compound, performance decreased compared to using Morgan Fingerprints for chemical representation (Fig. S8a,b).

Given the drug-like features of *ramA* inducers in our dataset, we applied our model to ∼12,000 DrugBank compounds^45^ (out-of-sample challenge, Methods), identifying 226 additional candidate inducers, the vast majority of which being human-targeted drugs (Fig. 6d). Most predicted inducers mapped to drug classes already represented in our dataset, primarily compounds targeting the nervous and respiratory systems (Figs. S8e, S9a). Experimental validation of 32 newly predicted compounds spanning diverse drug classes, chemical structures, and prediction scores confirmed 22 true inducers (Figs. 6e, S9a-c). This shows good predictive precision of our model (69%), an expressive increase compared to the hit rate in our initial screen (∼6.2% *versus* 1.07%), which is especially interesting for an out-of-sample challenge. A closer inspection of false positive predictions revealed that they more frequently contain molecular features negatively associated to *ramA* expression in our model compared to true positive examples. (Fig. S9d). Interestingly, we also noticed that *ramA* inducers in our dataset occupy a constrained molecular weight range, which was also not matched by falsely predicted candidates (Fig. 6f). This suggests that explicit inclusion of physicochemical properties could further improve model predictions. Nonetheless, our analyses revealed key molecular properties driving *ramA* induction, likely through RamR de-repression in *Salmonella*. Altogether, the identified and predicted RamA inducers in this study expand the previously reported space of *ramA*-inducing human-targeted drugs by ∼17-fold (Fig. 6g, Supplementary table 12), emphasizing a broad and previously unrecognized spectrum of human-targeted drugs that induce intrinsic antibiotic resistance genes in this pathogen.

## Discussion

Bacterial infections remain a major challenge for modern healthcare, requiring new strategies to counteract virulence and antibiotic resistance. For many genes involved in these processes, it remains unclear if and how their expression is affected by commonly used drugs, which can be critical in situations of polypharmacy. Here, we mapped a large network of compound-promoter interactions that captures transcriptional rewiring of virulence and antibiotic resistance associated pathways in *Salmonella*. We found a large set of compounds, in particular human-targeted-drugs, inducing transcriptional changes across different stress response pathways, the large majority of which were not previously described. We then integrated our chemically broad dataset with publicly available transcriptomic data, and revealed a new chemical inducer of PhoPQ-regulated genes, including a small RNA absent from our initial dataset. This example illustrates how this study can support mechanistic hypotheses beyond the initial setup of our screen, and prompted us to develop an interactive tool to facilitate data exploration by the community.

Our network exposes >200 non-antibiotic human-targeted drugs, several of them heavily prescribed and even over-the-counter medications, that modulate transcription of virulence or intrinsic antibiotic resistance genes in *Salmonella*. On the one hand, we uncovered human-targeted drugs with previously unknown impact on virulence, in particular through post-transcriptional regulation of SlyA. While the SlyA ligand binding properties have been mostly explored in the context of endogenous regulation of metabolism in *Salmonella*^35^, here we highlight SlyA as a potential mechanistic hub by which non-antibiotics can directly interfere with host-pathogen interactions. On the other hand, we show that human-targeted drugs trigger intrinsic antibiotic resistance in *Salmonella* by increasing efflux via RamA/RamR. There is increasing evidence suggesting that human-targeted drugs contribute to antimicrobial resistance, and that efflux may play a major role^65^. For instance, it has been shown that Enterobacteria are more resistant to human-targeted drugs due to multidrug efflux pumps^1,5^. Moreover, commensal-specific efflux modulation by human-targeted drugs has been shown to protect the gut microbiota from collateral damage by antibiotics^66^. We now show that human-targeted drugs can also directly and broadly increase the levels of multidrug efflux pumps in *Salmonella*, thereby increasing its resistance to antibiotics of different classes. Altogether, our findings suggest that human target drugs can influence the course of bacterial infections not only by modulating the microbiota as previously shown^1,2,5,6,67,68^, but also directly affecting virulence and intrinsic resistance pathways of a pathogen.

Many of the host-related molecules identified here have previously been associated with infection-relevant phenotypes, including altered bacterial growth, bioaccumulation in microbial communities, inhibition of efflux, and inhibition of *Salmonella* pathogenicity^1,2,28,30,68,69^ (Fig. S10a, b). However, in most instances, the molecular mechanisms underlying the described phenotypes are unknown, or are described for individual molecules^67–69^. Our approach enabled us to systematically discover molecular mechanisms of the impact of human-targeted drugs on virulence and intrinsic resistance, and revealed that *Salmonella* mounts remarkably uniform responses to diverse human-targeted drugs. While the perpetrators orchestrating these responses – RamA/RamR and SlyA – are reasonably well characterized ligand-responsive TetR- and MarR-family regulators, here we expand the chemical diversity of potential ligands far beyond what has been previously shown. This is the case of dantrolene, which is very different from the canonical salicylate-like molecules known to inhibit MarR-family regulators, including SlyA^35,60^. In fact, only ∼40% of the SlyA inhibitors and ∼25% of the *ramA* inducers identified in this study are structurally similar to previously reported MarR-family regulators ligands and RamA inducing chemicals^32,33,35,60,70,71^, respectively (Fig. S10c, d, Jaccard distance used as metric). This newly discovered chemical diversity, enabled us not only to rationalise chemical features of *ramA* inducers using machine learning, but also to predict >200 additional human-targeted drugs potentially increasing intrinsic antibiotic resistance in *Salmonella* with remarkable accuracy. Importantly, even though we acknowledge that our model might miss additional potential *ramA* inducers, it exposes an extensive and largely unexplored chemical space previously unknown to directly modulate bacterial regulatory networks.

TetR- and MarR-family regulators are widespread in bacteria and control a range of different biological functions for stress adaptation, including multidrug resistance, detoxification of xenobiotics, virulence, and metabolic adaptation^23–25,35,72^. Key questions to tackle in the future will be how generalizable our findings are across bacterial species. Here we highlight two scenarios: i) structural conservation of the regulators and lack of functional conservation of the regulons; and ii) poor conservation of the regulators and high functional overlap of the regulons. In the first scenario many TetR- and MarR-family regulators are indeed conserved, especially in closely related bacteria. Namely, RamR and SlyA are highly conserved among enterobacterial pathogens, including *Salmonella* Typhi and *Klebsiella pneumoniae*^72–74^. This leads us to speculate that the compounds uncovered in this study could target similar TetR- and MarR-family regulators across species, and indeed salicylate has been shown to interact with MarR-family regulators across several bacterial species including *E. coli*, *Salmonella*, and even *Mycobacterium tuberculosis*^75^. The regulons, however, are much more diverse, which makes direct generalization of phenotypes more challenging. For example, i) RamR controls antibiotic efflux in *Salmonella,* but outer membrane biogenesis in *Klebsiella*^72^; ii) SlyA regulates very distinct gene sets, even between closely related species such as *E. coli*, *Salmonella* and *Shigella*^73,74^, iii) here we showcase a very intricate cross-talk between MarR and SlyA regulation of the *marRAB* operon in *Salmonella*, which seems to overrule its canonical auto-regulation well-established in *E. coli*^39,76^. Therefore, we argue that the phenotype caused by each compound will most certainly change across species, but it is likely that the same compounds will have phenotypic consequences in other species if the regulators are conserved. For the second scenario – poor conservation of the regulators and high functional overlap of the regulons – we would like to highlight very recent results from collaborators, where we found that >100 human-targeted drugs increase expression of an efflux pump in *Campylobacter jejuni*, which is repressed by a structurally “RamR-distant” TetR regulator (manuscript in preparation). This striking conceptual and mechanistic overlap with our findings suggests that the mechanisms uncovered here may extend beyond *Salmonella* and represent a broader feature of pathogen response to host-associated environmental cues. Finally, we would like to highlight that our experimental and computational framework is easily generalisable to include other species, and can also be adapted to tackle different biological questions/systems, for example by exploration of different chemical spaces, other regulatory functions/genes/promoters, alternative representation of chemical features or modeling approaches.

## Acknowledgments

We thank Libera Lo Presti (University of Tübingen, GER) for providing feedback on the manuscript, the Brochado (University of Tübingen, GER) and Sharma lab (University of Würzburg, GER) members for discussions. We thank the Core Unit Systems Medicine (University of Würzburg, GER) for support with RNA sequencing, and the CSC-Finland for the computational resources.

## Author contributions

ARB, CLM, CMS and MKA conceived the study. MKA, AZ and ARB designed and optimized the high-throughput experiments. MKA performed the high-throughput experiments. MKA and ROA analysed the data. MKA, SB, LD, AB and EF designed and performed molecular cloning, cell culture and biochemical assays. TK and AR performed the ligand docking and molecular dynamic simulations. ROA, MKA, CLM and ARB implemented the machine learning model, including validation and interpretation. MKA and ARB wrote the manuscript. All authors discussed the results, edited and approved the final manuscript. ARB, CLM, CMS, TK and SW acquired funding and supervised this study.

## Data and Code Availability

An user-friendly interactive application to facilitate navigation of our dataset by the scientific community is available at (link). Trajectory and interaction data are publicly available in the Zenodo repository (DOIs: 10.5281/zenodo.15582334 and 10.5281/zenodo.17858054). The data supporting the findings of this study will be available as source data & supplementary files/Github. All code will be made available on GitHub.

## Supplementary figure legends

**Figure S1.**
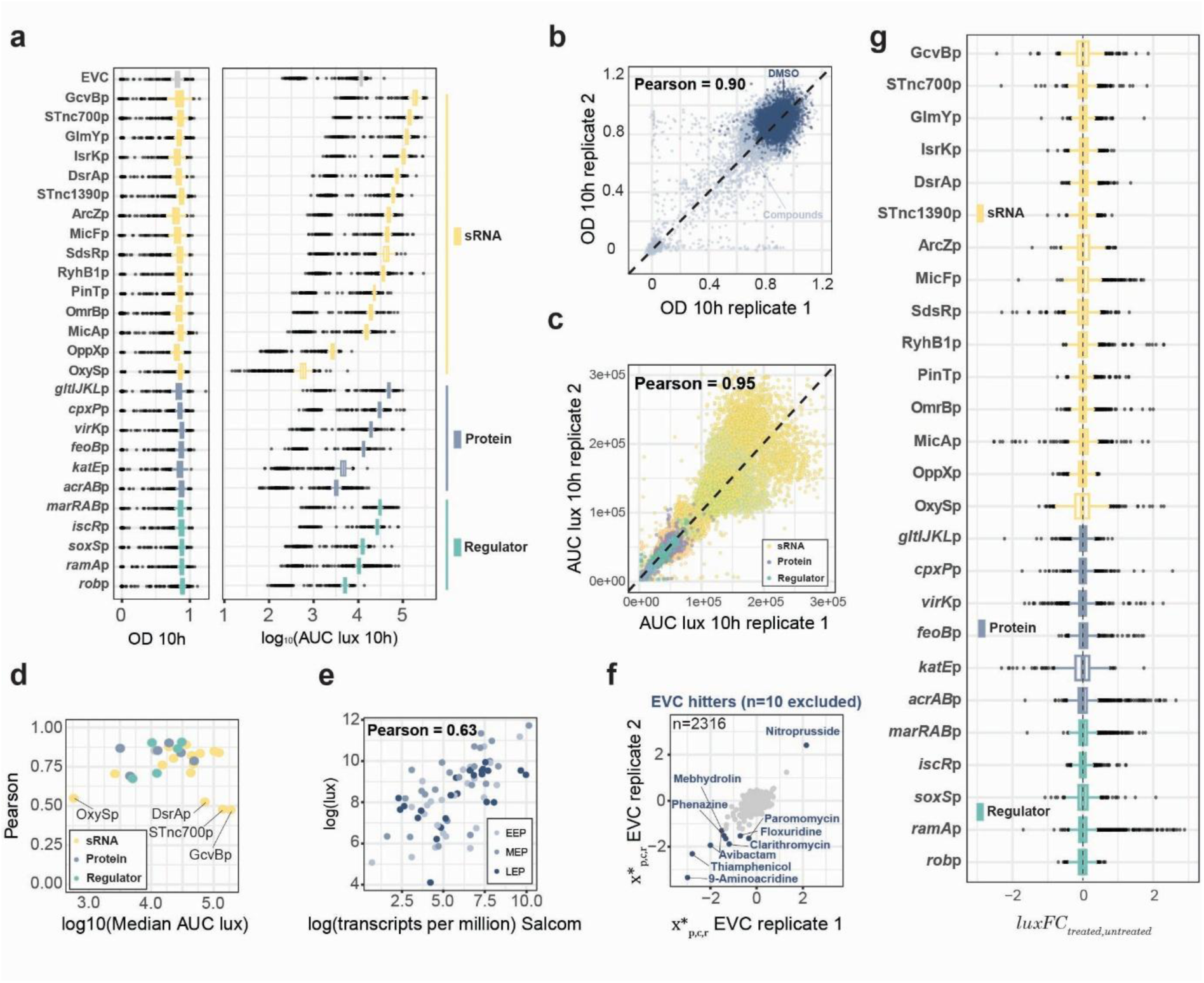
Quality control and data analysis. **a)** Promoter library has a wide dynamic range of reporter activity, and does not influence *Salmonella* fitness. Growth (OD at 10h incubation, left) and reporter activity (log_10_(AUC of luminescence-over-time up until 10 h, right) of all data. Boxplots refer to growth and luminescence data of 2415 compounds as the average between the two biological replicates. Boxplots indicate 25th, 50th and 75th percentiles, and whiskers extend up to 1.5 times the interquartile range (IQR) from the 25th and 75th percentiles (applies to **a** and **g**). **b and c)** Pearson correlation of growth **(b)** and promoter activity **(c)** between the two biological replicates for all 62,790 compound-promoter pairs tested. In **b**, wells containing the vehicle control (DMSO) are depicted in blue, and those containing the compounds are depicted in grey. **d)** Low- and highly-expressed promoters are overall noisier. Scatterplot shows the promoter-specific Pearson replicate correlations of AUC luminescence until 10h against the promoter activity levels (median AUC luminescence until 10h). Promoters that present Pearson <0.6 are highlighted. **e)** Luminescence activity of the promoter library correlates with transcript levels from an independent study ^13^. Transcript levels (tpm) of the genes under control of the promoters of our reporter library were extracted from the SalCom database ^13^. Scatterplot shows the comparison of luminescence (this study) x transcript levels (SalCom) for distinct growth phases: EEP (early exponential phase, OD_600nm_ = 0.1), MEP (mid exponential phase, OD_600nm_ = 0.3) and LEP (late exponential phase, OD_600nm_ = 1.0). **f)** Effect of the screened compounds on the empty vector control. Scatterplot shows the median-centered, growth-normalized luminescence (Methods) of the empty vector control (EVC) in response to 2316 compounds that did not inhibit the growth of *Salmonella*. The ten compounds that were not considered for further analysis by substantially interfering with the EVC (growth-normalized luminescence ≥1.5 log_2_ fold-change for at least one replicate) are indicated. **g)** Boxplots showing the promoter activity (*luxFC_treated_*_/*untreated*_) per promoter across all compounds retained after quality control (n=2306).

**Figure S2.**
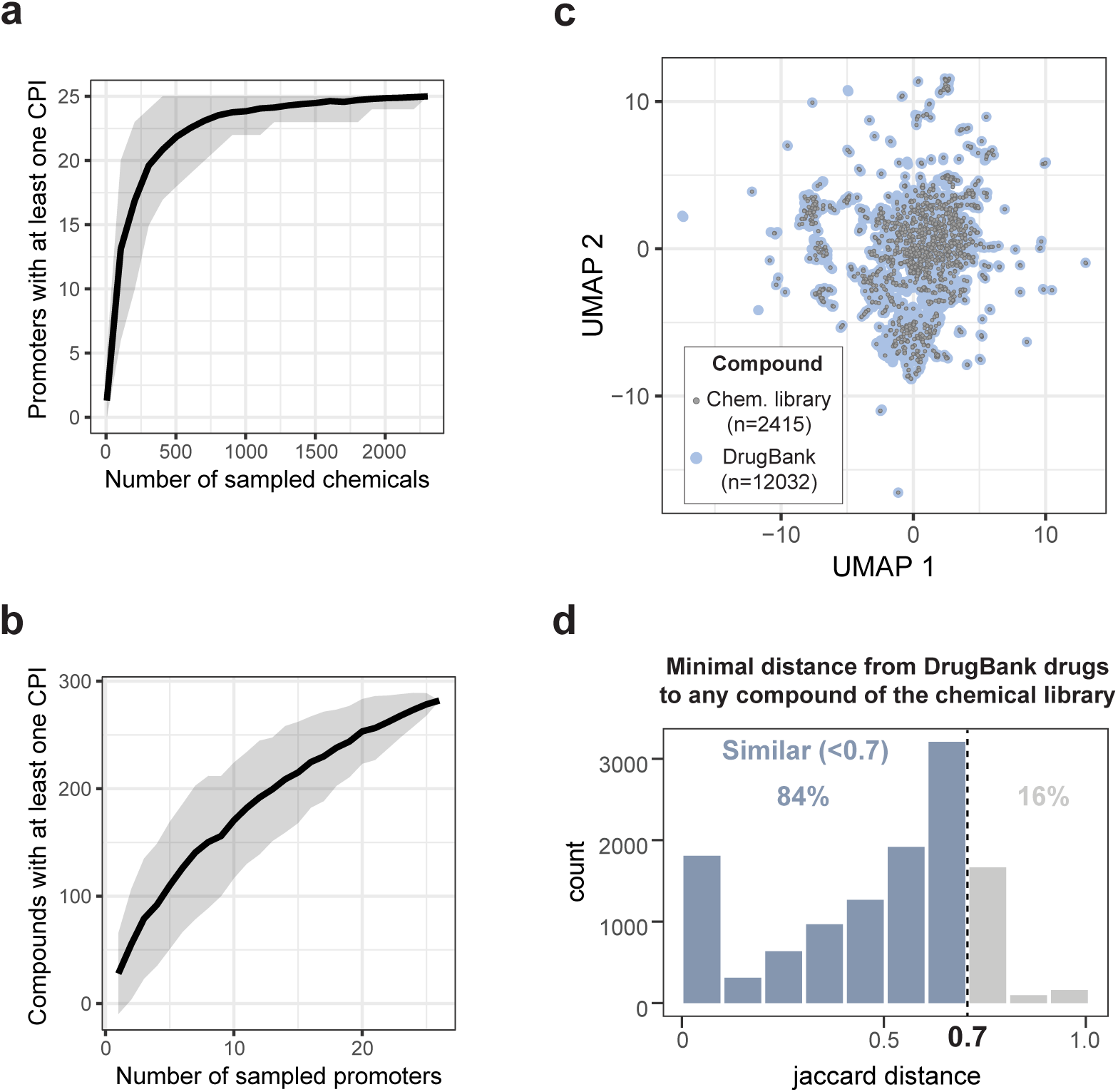
Sampling depth of the high-throughput screening approach. **a)** Number of compounds saturates CPI discovery within the selected promoters. Curves show the dataset rarefaction analysis of the number of promoters with at least one CPI per number of compounds screened. The tick line represents the mean and the shaded areas the standard deviation across 100 random sampling interactions (Methods). **b)** Number of screened promoters did not saturate CPI discovery. Curves show the dataset rarefaction analysis of the number of compounds with at least one CPI per number of screened promoters. Tick lines represent the mean and the shaded areas the standard deviation across 100 random sampling interactions (Methods). **c)** Visualization of the chemical space coverage of the compound library used in this study in comparison with the DrugBank; Morgan fingerprints were used to create an Uniform Manifold Approximation and Projection (UMAP) overlaying the compound library onto the Drugbank compounds. n indicates the number of compounds. **d)** Compound library used in this study reflects the chemical diversity of the Drugbank. Histogram shows the number of compounds from the Drugbank distributed according to their Jaccard distance to the closest compound from our library. Assuming a threshold of 0.7 - conventionally considered as an indicator of similarity - 84% of all DrugBank molecules have a similar chemical represented in our compound library.

**Figure S3.**
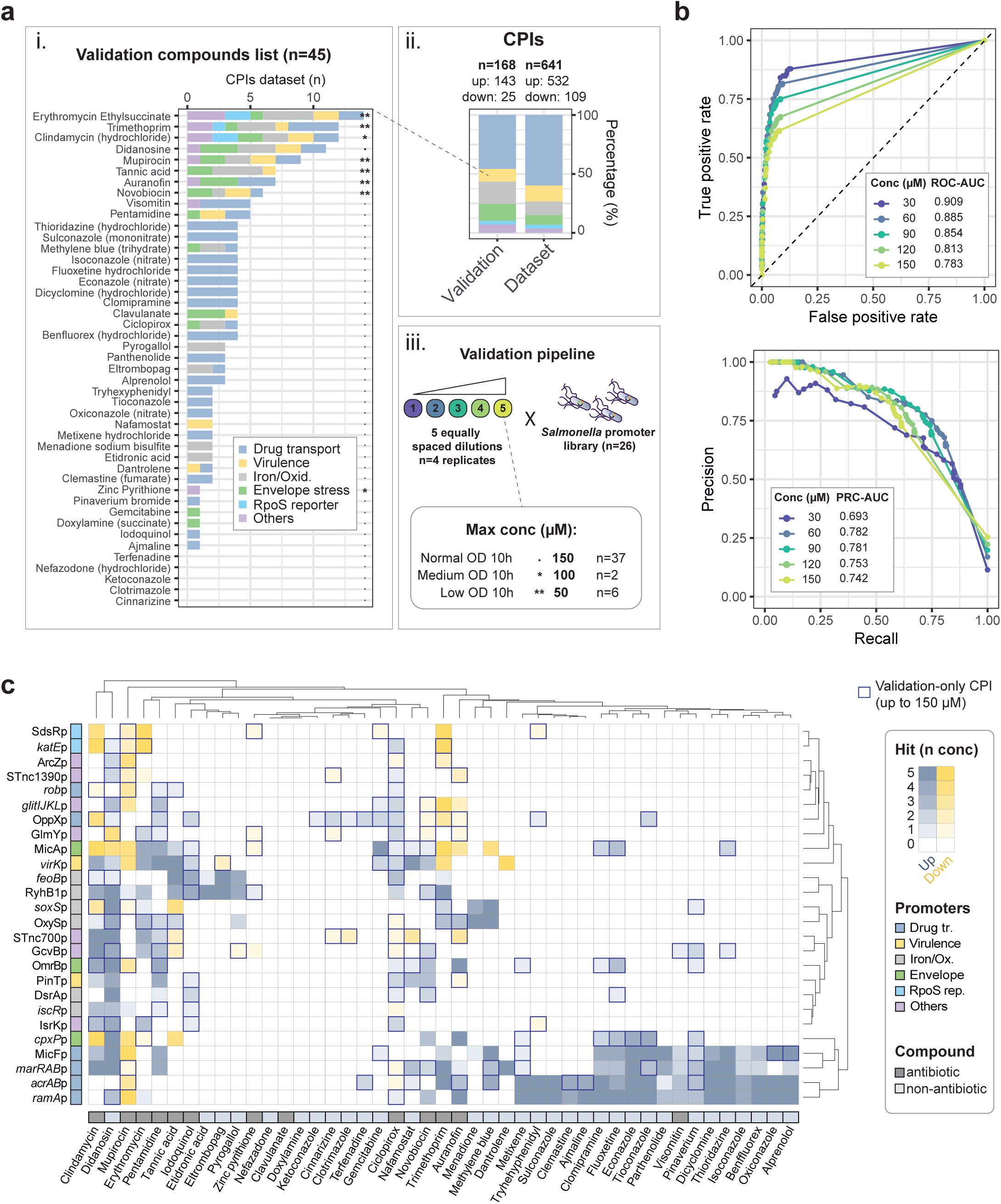
Concentration-resolved validation. **a)** Schematic representation of the systematic concentration-resolved validation of the primary screen; i) 45 compounds selected for validation. Bars show the number of CPIs and the stress pathway of the promoters as observed in the primary screen; ii) The validation subset represents a good sampling of the CPIs diversity of the primary screen. Bars show the proportion of stress pathways of the promoters involved in the 168 CPIs selected for validation, as well as that of the 641 CPIs obtained in the dataset (primary screen); iii) Experimental pipeline of the validation. Briefly, 45 compounds were tested against the promoter reporter library at five equally spaced concentrations (Supplementary table 7, Methods). The maximum concentration was either 150 µM (n=37), 100 µM (n=2) or 50 µM (n=6), depending on compound toxicity towards *Salmonella*, and it is indicated by symbols (dot - 150 µM, one asterisk - 100 µM and two asterisks - 50 µM). **b)** Performance metrics of the dataset validation. Receiver Operating Characteristic (ROC, top) and precision-recall (PRC, bottom) curves of the primary screen results compared with concentration-resolved screen validation (gold-standard), done for each individual concentration (30 to 150 µM). **c)** Overview of the validation results. The heatmap shows the results of the 1170 compound-promoter pairs tested for validation. The intensity of the tile refers to the number of concentrations that resulted in CPIs in the validation set (Methods), and color refers to the direction of the interaction. CPIs that were only observed in the screen validation and not in the large screen (false negatives), are highlighted with borders.

**Figure S4.**
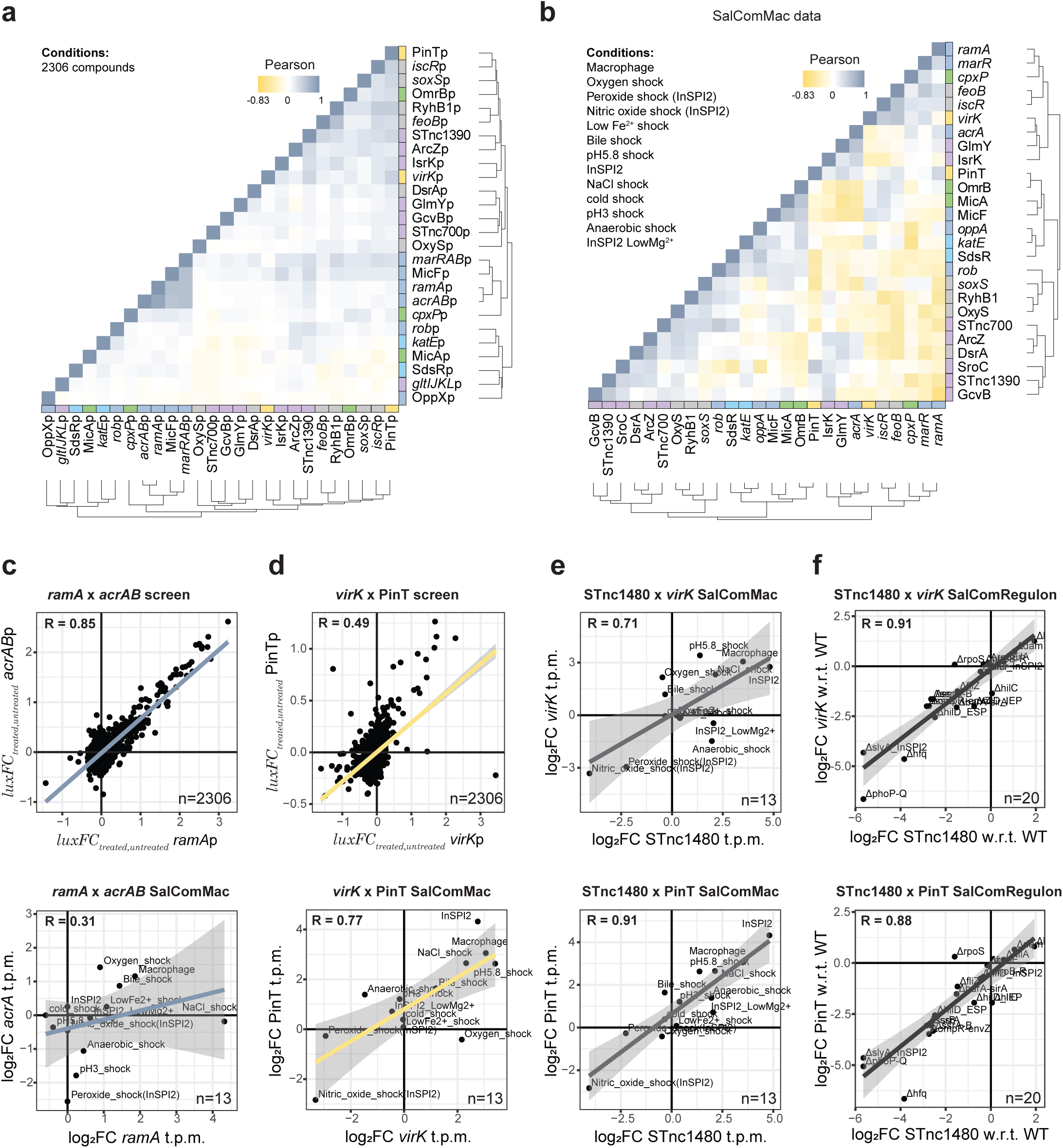
Benchmarking the results of this study against a previously published dataset - SalComMac. **a-b)** Heatmaps showing Pearson correlation coefficients between all promoter pairs from our dataset **(a)** and Pearson correlation coefficients between gene expression profiles across 13 infection-relevant conditions from SalComMac **(b)** ^44^. Only the genes corresponding to the 26 promoters included in our screen are represented. Pearson correlation coefficients between all promoter pairs from our dataset were calculated using promoter activity (*luxFC_treated_*_/*untreated*_) across 2306 compounds (that passed quality control). Pearson correlation coefficients between the 26 genes from SalComMac were calculated using transcript fold-change across 13 infection-relevant conditions **c-d)** Showcasing specific examples for benchmarking our dataset against SalComMac. Scatterplots with correlation of promoter activity/gene expression between two different promoters/genes across all compounds tested here or conditions from SalComMac ^44^. n indicates the number of compounds/conditions. **c)** *ramA*p x *acrAB*p are strongly correlated in our screen (top) but weakly correlated in SalComMac (bottom). **d)** *virK*p and PinT are strongly correlated in both our screen and SalComMac. **e)** Expression of the sRNA STnc1480 is strongly correlated with *virK* and PinT in SalComMac. Scatterplots showing the correlation of expression between *virK* (top) or PinT (bottom) and STnc1480 across all conditions from SalComMac ^44^. **f)** STnc1480 is under the same regulatory control of *virK* and PinT. Scatterplots showing the correlation of expression between *virK* (top) or PinT (bottom) and STnc1480 across *Salmonella* knockout strains of major stress regulators from <u>SalComRegulon</u> ^46^. log_2_FC w.r.t. WT refers to the transcript levels in the indicated regulator knockout mutants in relation with the WT strain. For **c-f**: R represents Pearson correlation of the data shown, and lines refer to a fitted linear model of the data shown, with 95% confidence intervals depicted by grey shades.

**Figure S5.**
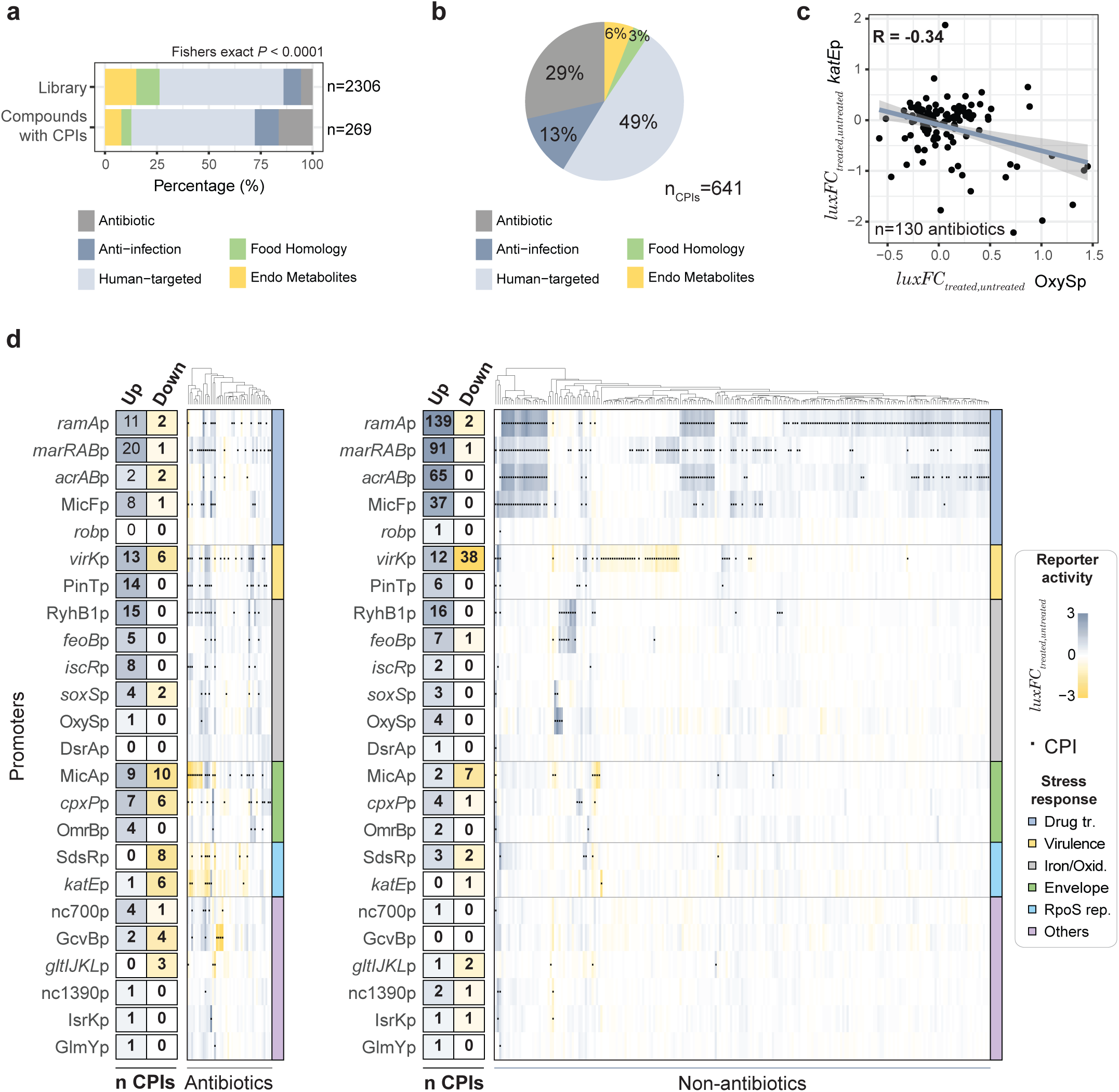
Antibiotics and non-antibiotic compounds elicit distinct regulatory responses in *Salmonella* (extended information). **a)** CPIs are enriched for antibiotics. Bars show the proportion of the compounds in the whole library (top, after quality control, n=2306), and involved in at least one CPI (bottom, n=269), distributed by the respective compound classes. P represents the p-value of a Fisher’s exact test (“greater than”) comparing the number of antibiotics versus the number of non-antibiotics (all other classes together), depending on whether they are or not involved in at least one CPI (total nr compounds = 2306). **b)** Human-targeted drugs are involved with nearly half of the CPIs. Pie chart showing the distribution of CPIs according to compound class. n indicates the total number of CPIs. **c)** OxyR-regulated promoter OxySp and RpoS-regulated promoter *katE*p are anticorrelated in response to antibiotics. Scatterplot shows the correlation of promoter activity (*luxFC_treated_*_/*untreated*_) between OxySp and *katE*p across all antibiotics in our screen after quality control. n indicates the number of antibiotics. R represents Pearson correlation of the data shown, and lines refer to a fitted linear model of the data shown, with 95% confidence intervals depicted by grey shades. Heatmaps depict promoter activity (*luxFC_treated_*_/*untreated*_) of all promoters with at least one CPI across 44 antibiotics (left) and 225 non-antibiotics (right) with at least one CPI. Tables show the number of CPIs per promoter among the represented compounds (antibiotics on left and non-antibiotics on the right side). Significant CPIs are highlighted by black dots. The dendrograms show hierarchical clustering of the compounds across promoters based on Euclidean distances with complete linkage.

**Figure S6.**
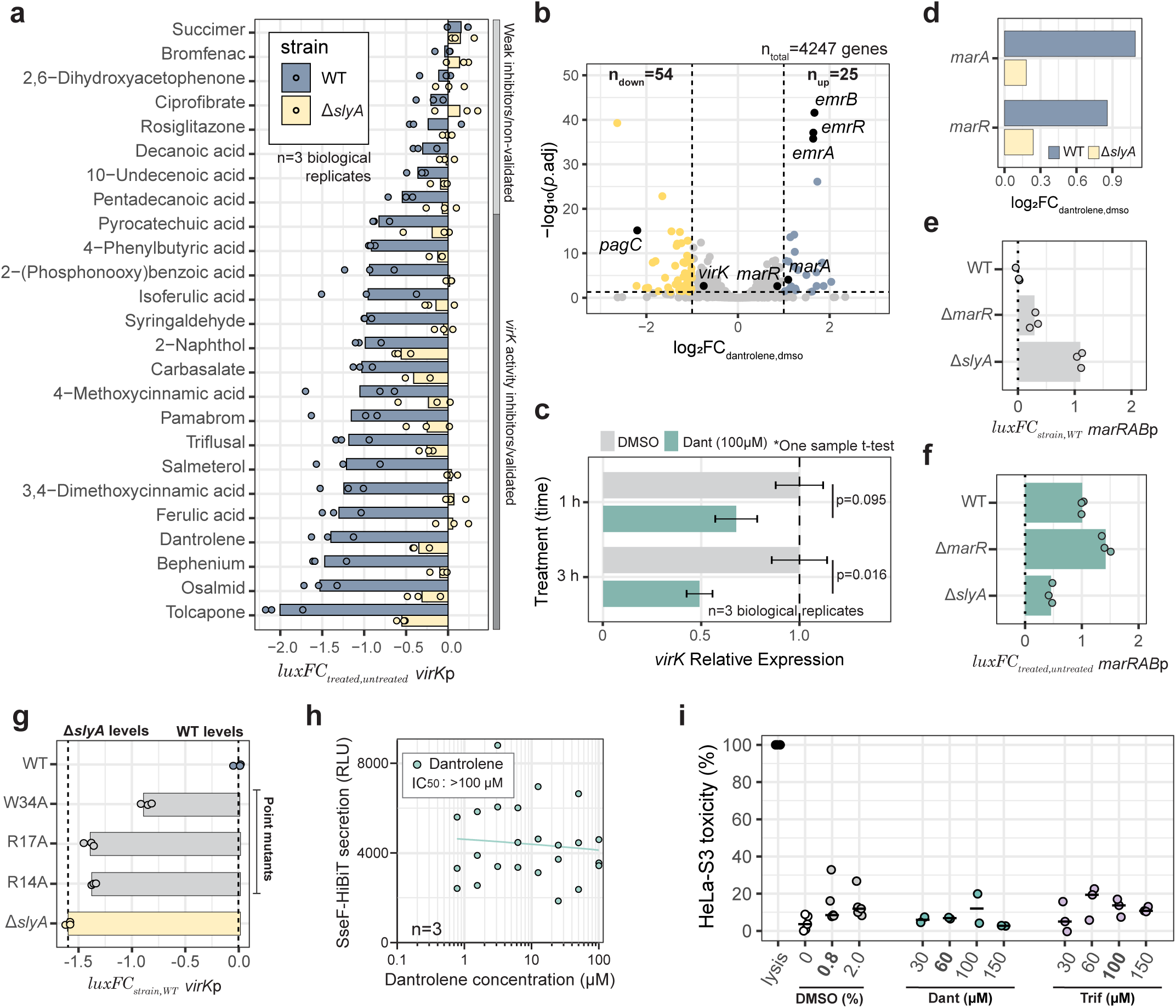
Virulence regulator SlyA is regulated by non-antibiotic compounds (extended). **a)** Non-antibiotics decrease *virK*p promoter activity via *slyA*. Bars show the average change in *virK*p promoter activity (*luxFC_treated_*_/*untreated*_) in response to 25 non-antibiotics (100 µM). Data is shown for wild-type (blue) and Δ*slyA* (yellow) deleted *Salmonella*. n indicates the number of biological replicates. **b)** The impact of dantrolene on the transcriptional landscape of wild-type *Salmonella* recapitulates typical SlyA-, MarR- and EmrR-mediated responses. The volcano plot shows the RNA-sequencing results based on three biological replicates after DESeq2 analysis. Overnight cultures of *Salmonella* were diluted to OD_600 nm_ ∼0.01 in LB, grown until OD_600 nm_ ∼0.1, and dantrolene (100 µM) or DMSO were added for 1 hour before RNA extraction. Significantly down- and up-regulated genes are marked in yellow and blue, respectively, and the corresponding count is indicated by n_up_ and n_down_. Genes previously reported to be part of SlyA (*pagC* and *virK*), MarR (*marA* and *marR*) or EmrR (*emrA*, *emrB* and *emrR*) regulations are highlighted (positive controls). n_total_ indicated the total number of genes detected. The dotted vertical and horizontal lines show the thresholds used for statistical analysis (Methods). **c)** RT-qPCR confirms dantrolene-mediated down-regulation of *virK*. Bars show the transcriptional change of *virK* in response to dantrolene (100 µM) in comparison to the vehicle (DMSO). Treatment was done for 1 or 3 hours and *rpoS* was used as reference (Methods). p-values were calculated using one-sample t-tests against a theoretical mean of 1 (no change from control). The bars and error-bars show average and standard deviation across n biological replicates, respectively. **d)** Deletion of *slyA* impairs dantrolene-mediated induction of *marA* and *marR*. Barplot shows the changes in transcript levels of *marR* and *marA* in wild type and Δ*slyA Salmonella* upon dantrolene treatment (log_2_FC_dantrolene,DMSO_ from the RNA sequencing experiment). The average of three biological replicates is shown. **e)** Deletion of *slyA* influences the basal expression levels of *marRAB*p activity. Barplots show the average change of *marRAB*p promoter activity (reporter-based, *luxFC_strain_*_,*WT*_) of untreated *Salmonella* mutants (Δ*slyA* and Δ*marR*) in comparison to the wild type. Bars show the average of three biological replicates. **f)** Dantrolene-mediated induction of *marRAB* promoter activity is *slyA*-dependent. Barplots show the average change of *marRAB*p promoter activity (reporter-based, *luxFC_treated_*_/*untreated*_) upon dantrolene treatment (100 µM) for three genetic backgrounds: wild type, Δ*slyA* and Δ*marR* deleted *Salmonella*. Bars show the average of three biological replicates. **g)** Effects of *slyA* point mutations on its regulatory activity towards *virK*p. Barplots show the average change of *virK*p promoter activity (reporter-based, *luxFC_strain_*_,*WT*_) of untreated *Salmonella* carrying different SlyA variants in comparison to the wild type. Dashed lines show the average change of *virK*p promoter activity for the wild-type (reference) and Δ*slyA* (far left, negative control). Bars show the average of three biological replicates. **h)** SPI-2 secretion activity upon dantrolene treatment. A curve of SPI-2 secretion activity over increasing dantrolene concentration is shown. Secretion activity was measured as previously established ^62^. n indicates the number of biological replicates. Results are inconclusive, most likely due to dantrolene degradation in assay conditions. **i)** Dantrolene and triflusal did not show cytotoxic effects in HeLa-S3 cells. Dotplot shows HeLA-S3 viability as measured by LDH release assay (Methods). Each point corresponds to individual biological replicates as the average of three technical replicates. Data shown based on 2-5 biological replicates. Toxicity levels below 20% are generally considered low.

**Figure S7.**
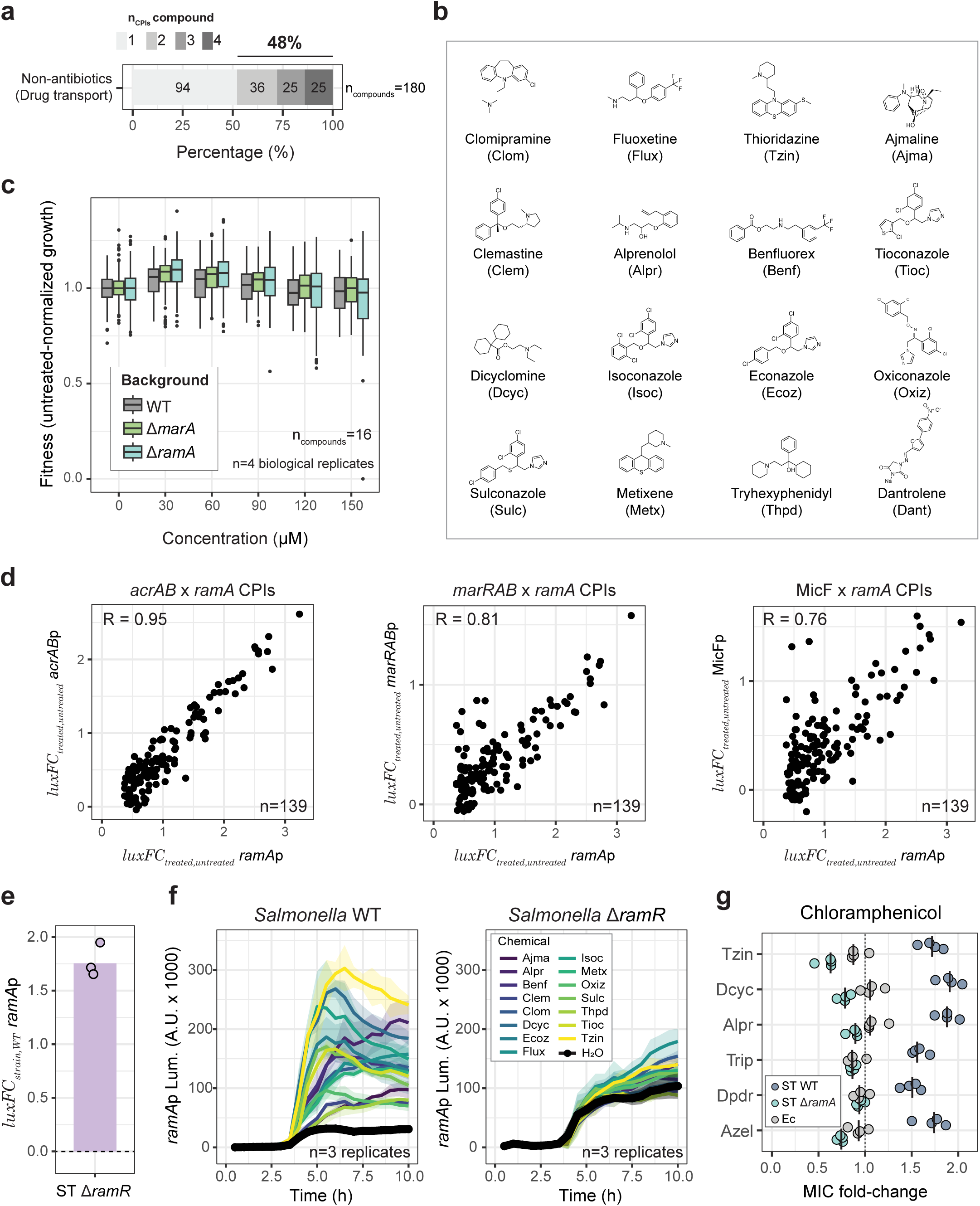
Extended information on *ramA*-mediated regulation by non-antibiotics. **a)** Extensive cross-regulation of drug-transport associated promoters by non-antibiotics. Bar showing the percentage of non-antibiotics that are involved in 1, 2, 3 or 4 CPIs with drug-transport associated promoters. Numbers inside the bar represent the exact number of compounds with the color-coded corresponding number of CPIs. **b)** Compounds that change activity of drug-transport associated promoters have diverse chemical structures. Chemical structures of the non-antibiotics used in Figure 5 b&d. **c)** Non-antibiotics do not affect *Salmonella* growth in Δ*ramA* or Δ*marA* deletion backgrounds. Boxplots show the fitness of *Salmonella* wild type and Δ*ramA* or Δ*marA* deletion mutants upon treatment with the 16 compounds that change activity of drug-transport associated promoters tested in Fig. 5 b and d, and shown in Figure S7b. The data shown contains the (pooled) fitness data of the EVC and the four drug-transport associated promoter reporter strains across 4 biological replicates, for each individual genetic background and specific concentration (each boxplot is derived from 320 data points). Fitness refers to AUC of the growth curve (OD_600 nm_ up until 10h) of treated divided by pooled untreated *Salmonella* (all individual genetic background handled separately). Boxplots indicate 25th, 50th and 75th percentiles, and whiskers extend up to 1.5 times the interquartile range (IQR) from the 25th and 75th percentiles. **d)** Promoter activity of drug-associated promoters correlates with that of *ramA*p. Scatterplot shows the correlation of promoter activity (*luxFC_treated_*_/*untreated*_) between *acrAB*p (left), *marRAB*p (middle) or MicFp (left) and *ramA*p across the 139 non-antibiotics involved in a CPI with *ramA (n)*. R represents Pearson correlation of the data shown. **e)** Deletion of *ramR* confirms increased basal expression of *ramA*p in the reporter system. Barplot shows the average change of *ramA*p promoter activity (reporter-based, *luxFC_strain_*_,*WT*_) of untreated *Salmonella* Δ*ramR* in comparison to the wild type. Bar shows the average of three biological replicates. **f)** Deletion of *ramR* impairs increase in *ramA*p promoter activity elicited by non-antibiotics. Raw luminescence curves of *ramA*p expression for the fifteen compounds from Fig. 5d for *Salmonella* wild-type (left) and Δ*ramR* (right). Lines represent the average and the shaded area the 95% confidence intervals of three biological replicates. Untreated controls (H_2_O) indicate the basal expression levels of each genetic background. **g)** *ramA* inducers impact the minimal inhibitory concentration of AcrAB-TolC substrate chloramphenicol. Dotblots showing ciprofloxacin MIC fold change caused by the 6 *ramA* inducers from Fig. 5f for *Salmonella* (ST) wild type *ΔramA*, as well as for *E. coli* (Ec). Four biological replicates are shown.

**Figure S8.**
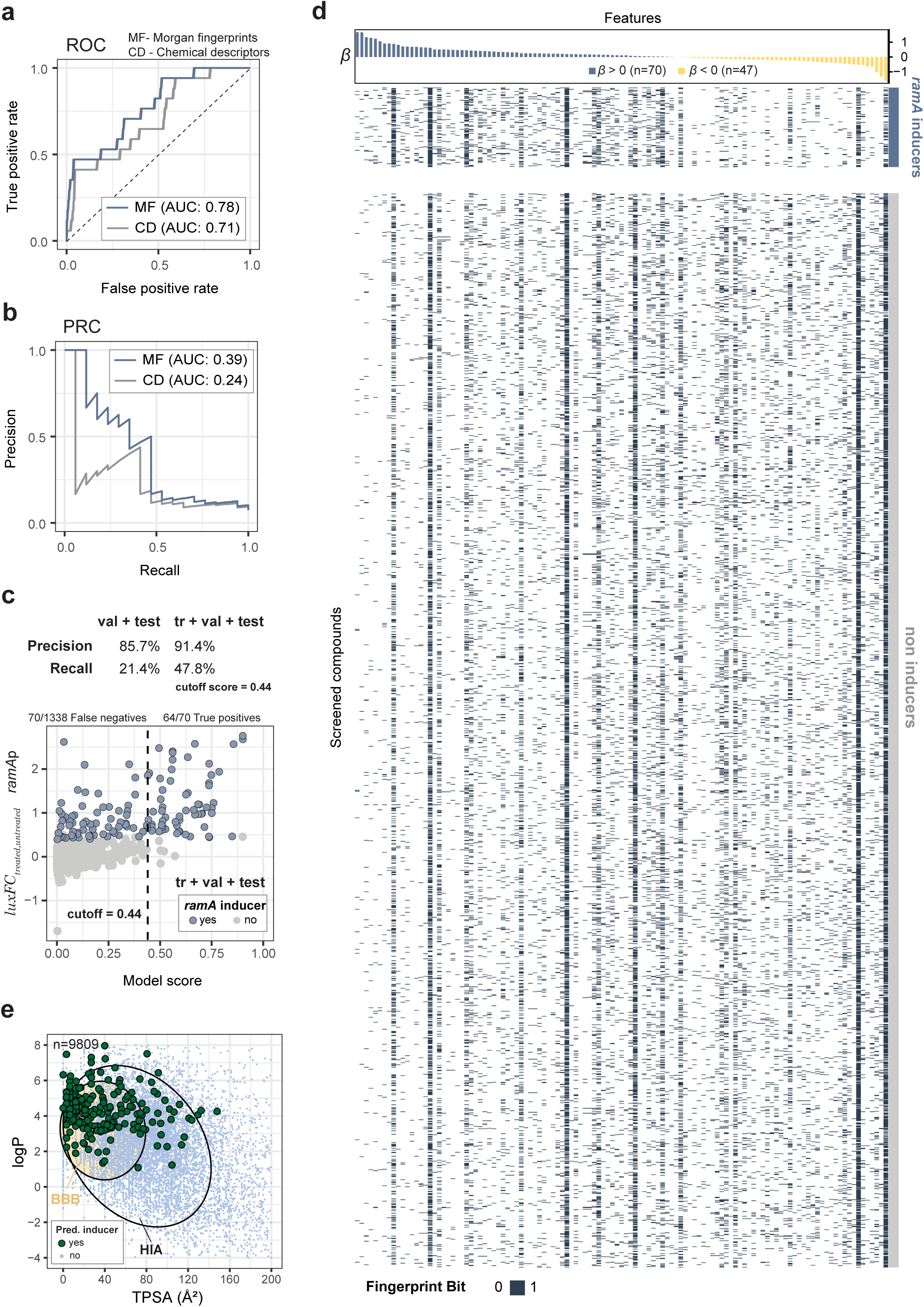
Analysis of molecular features and chemical properties of *ramA* inducers. **a and b)** Model performance metrics. Receiver Operating Characteristic (ROC, a) and precision-recall (PRC, b) curves for the LASSO Logistic regression models for prediction of *ramA* inducers. Curves corresponding to the LASSO logistic regressions based on Morgan fingerprints (blue) and chemical descriptors (gray) are shown. The area under the curves (AUC) shown were estimated for the validation set. **c)** Predictive model shows high precision and limited sensitivity. Scatterplot shows experimentally measured *ramA*p promoter activity and the corresponding score obtained with the LASSO Logistic regression model based on Morgan fingerprints. True positives and false negatives were called based on the cutoff of 0.44 on model scores. The table on the top shows precision and recall for the validation+test sets (val + test) or validation+test+training set (tr + val + test) when the cutoff of 0.44 is used on model scores. **d)** Morgan fingerprint bitmaps of all compounds in the large screen. Top: Barplot showing the model coefficients (*β*) for the 114 non-zero molecular features from the LASSO logistic regression. Bottom: bitmap showing the presence/absence of each one of the 114 features in the 2306 compounds used in our large screen, classified as *ramA*p inducer (top) or non-inducer (bottom). **e)** Model-predicted *ramA* inducers mostly correspond to drugs that cross the brain-blood barrier. BOILED egg plot of the compounds from DrugBank, represented by a scatterplot of logP (partition coefficient) and TPSA (topological polar surface area). The highlighted areas represent typical logP and TPSA ranges of compounds absorbed in the human intestine (human intestine absorption, HIA, white) and compounds that cross the blood-brain barrier (BBB, yellow). Model-predicted *ramA* inducers are highlighted in dark green.

**Figure S9.**
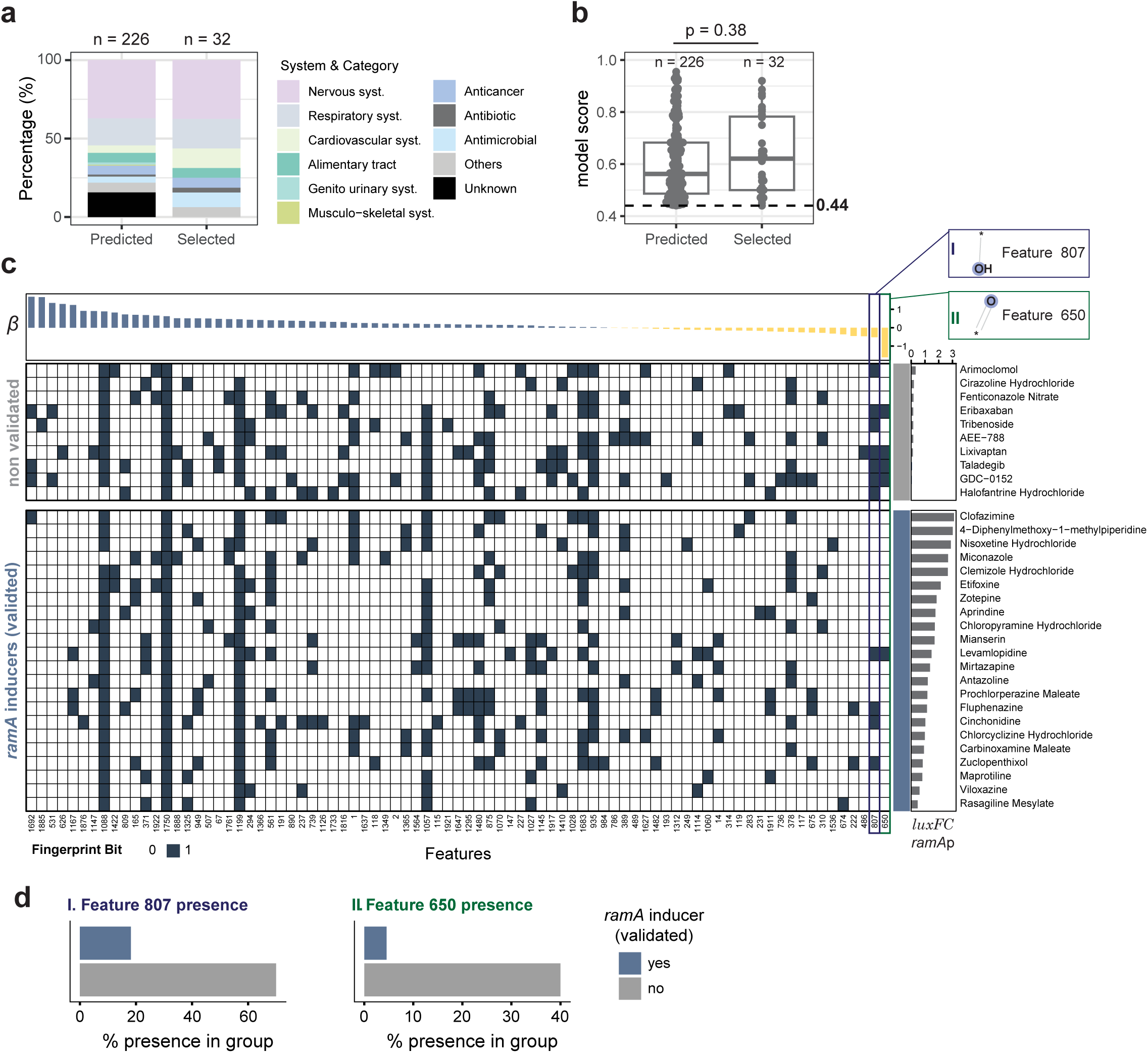
Out of sample challenge of the *ramA* inducers predictive model. **a)** The 32 candidate *ramA* inducers selected for experimental validation of the predictive model cover most drug categories represented in all newly predicted *ramA* inducers. Bars show the proportions of predicted *ramA* inducers distributed according to drug categories. The left bar shows all predicted new inducers from the DrugBank, while the right bar focuses on the 32 selected drugs for experimental validation. **b)** The 32 candidate *ramA* inducers selected for experimental validation of the predictive model cover a broad range of model scores. Boxplots show the distribution of model scores for the 226 predicted and the subset of 32 *ramA* inducers selected for experimental validation. The individual model scores are shown as overlaid dots. The dashed line represents the model score cutoff used to classify predicted *ramA* inducers (Methods). The p-value of a double sided Wilcoxon rank-sum test is shown. **c)** Non-validated compounds have more model features with negative coefficients. Morgan fingerprint bitmaps of the 32 compounds used for experimental validation of new predicted *ramA* inducers. Top: Barplot showing the model coefficients (*β*) for the non-zero molecular features across the 32 compounds. Bottom: bitmap showing the presence/absence of each one of the non-zero-coefficient features in the 32 compounds, classified as *ramA*p validated inducer (bottom) or not-validated (top) inducer. The strong negative features are highlighted. Experimental validation of increased *ramA*p promoter activity is shown as average *luxFC_treated_*_/*untreated*_ across 2-3 biological replicates on the right side of the bitmap. **d)** Barplots showing the percentage tested 32 compounds containing “negative features” 807 (left) and 650 (right), grouped as validated (blue) and non-validated (gray) *ramA* inducers.

**Figure S10.**
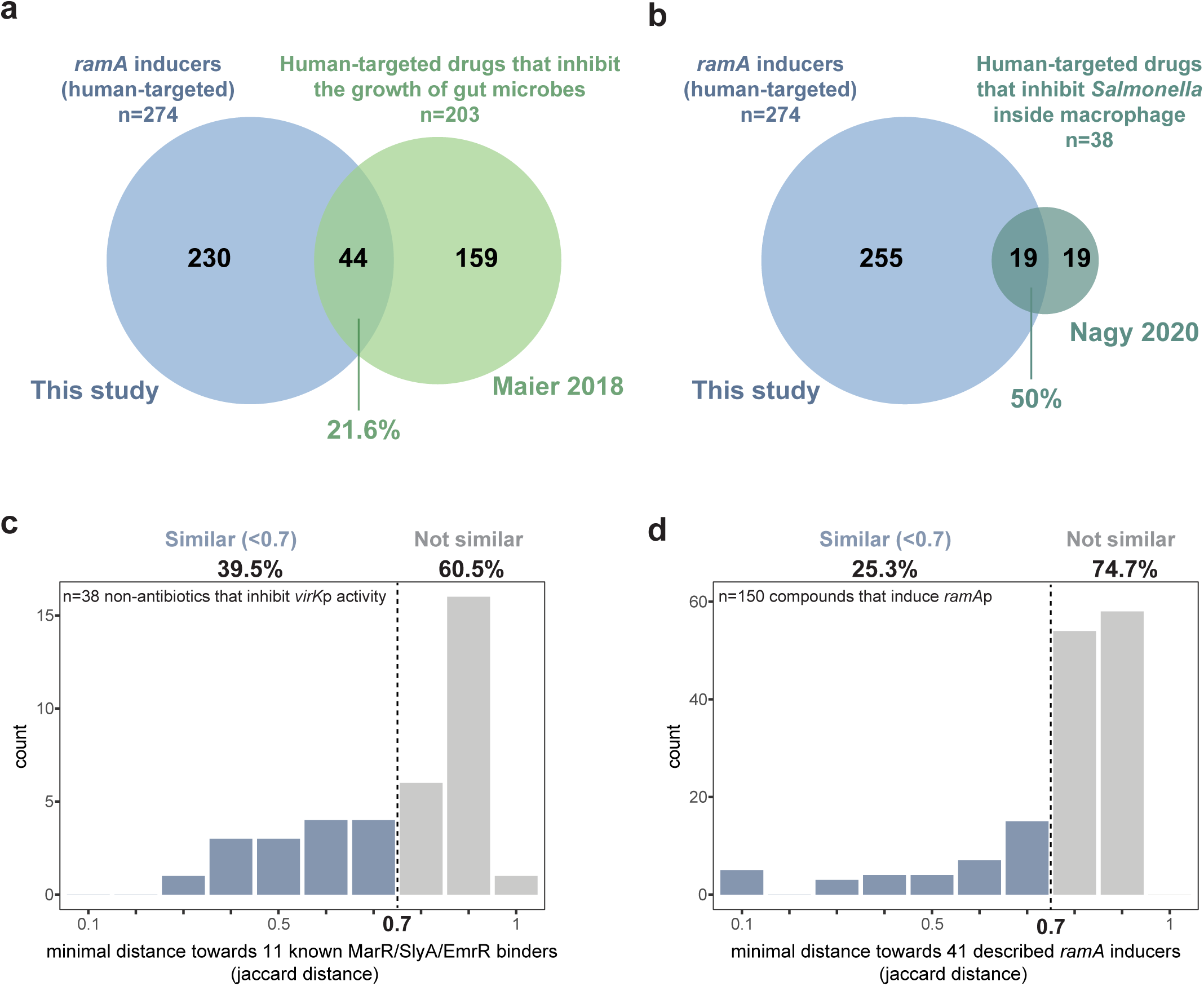
Extensive chemical diversity underlies drug-mediated modulation of bacterial regulatory networks. **a)** Intersection of *ramA* inducers with human-targeted drugs that impact growth of gut microbes ^1^. **b)** Intersection of *ramA* inducers with human-targeted drugs previously shown to inhibit *Salmonella* during macrophage infection ^4^. Percentages shown are calculated based on the number hits of the compared dataset in both cases (green). All human-targeted drugs from all datasets were taken into account (and not other compounds such as antibiotics, anti-infectives or metabolites), independently of whether the same drugs were tested in all studies. **c)** This study increases the chemical diversity of ligand-based SlyA regulation. Histogram shows the number of non-antibiotics that decrease *virK*p promoter activity distributed according to their Jaccard distance to the closest compound within a set of 11 compounds known to bind to MarR-family regulators (Supplementary table 13). **d)** This study increases the chemical diversity of compounds previously shown to induce *ramA*. Histogram shows the number of compounds that increase ramAp promoter activity (large screen data only) distributed according to their Jaccard distance to the closest compound within a set of 41 compounds known to increase *ramA* expression (Supplementary table 13). In **c** and **d** a threshold of 0.7 was considered as an indicator of similarity.

## Methods

### Culture conditions, reporter plasmid and strain construction

*Salmonella enterica subsp. enterica ser.* Typhimurium 14028S (*Salmonella*) was used as wild type, template and host for the reporter library and knockout mutant generation (NCBI Nucleotide Genbank entry CP001363). *Salmonella enterica subsp. enterica ser.* Typhimurium NCTC 12023 was used for the SPI-2-encoded Type III secretion system activity assay, and *Salmonella enterica subsp. enterica ser.* Typhimurium SL1344 was used HeLa-S3 infection assays. Bacteria were generally grown in Lysogeny Broth (LB) Lennox adjusted to pH 7.5 at 37 °C, unless stated otherwise. Medium was supplemented with carbenicillin (50 µg/mL, CatNo Cay20871-5, BioMol) or kanamycin (50 μg/ml, CatNo. K1876-5G, Sigma-Aldrich-Merck) when selection pressure was required.

Benchling was used for plasmid and primer design. Plasmid pASCoT-Lux was used as backbone for the promoter reporter library, where generally 500 bp upstream of the start-codon of the target gene were cloned upstream of the reporter operon *luxCDABE*. The pASCoT-Lux was constructed from pTU175-oriT-Lux ^38^, where the spectinomycin resistance cassette was replaced by the ampicillin resistance gene from pET21a(+). The vector contains a single XhoI restriction site 37 bp upstream of *luxC* (where a ribosome binding site is located), which was used to linearize the plasmid, and clone the promoter regions using Gibson assembly ^77^. For promoters containing 5’-UTR (OppX) or 3’-UTR sRNAs (SroC), the TSS of the primary transcript was considered ^13^. Promoter PCR amplification was performed with Q5 polymerase following supplier instructions (CatNo. M0491S, New England Biolabs (NEB), USA). Plasmid transformation was done by electroporation ^78^.

Δ*phoP, ΔphoQ, ΔslyA,* Δ*ramA* and Δ*ramR* were derived from a non-essential knockout library ^79^. The presence of the kanamycin resistance cassette in gene loci was confirmed using PCR, and then back transduced into wild type using P22. Plasmid pCP20 was used for kanamycin resistance cassette removal. Scarless deletions of *marA* and *marR*, as well as *slyA* point mutations (R14A, R17A and W34A), were done though two-step allelic exchange method using the suicide vector pNTPS138-R6K as previously described, using kanamycin 50µg/mL for selection, and 10% sucrose (wt/vol) for counter-selection ^80^. Eco-RV (CatNo. R0195S, New England Biolabs (NEB), USA) was used for plasmid linearization, and 500-750 base-pair fragments used for homologous recombination were inserted with Gibson assembly ^77^. DNA transfer to *Salmonella* was done by conjugation using *E.coli* WM3064.

All plasmids inserts and genomic modifications were confirmed with Sanger sequencing. All primers and plasmids used in this study are listed in Supplementary tables 14 and 15, respectively. All strains (Supplementary table 16) and plasmids constructed during this study are available from the authors upon request.

### Compound-promoter combinations screen

High-throughput compound mediated transcriptional response of the target promoters (n=26 plus the promoter-less **e**mpty **v**ector **c**ontrol [EVC], Supplementary table 1) to the 2,415 individual compounds (Supplementary table 2) was done in an liquid handling automated workstation (Beckman Coulter, USA), with a Biomek i7 liquid handler, a Cytomat2 shaking incubator (Thermo Scientific, USA), and a plate reader Synergy H1 BioTek with Gen5 3.09 (Agilent, USA), as previously described ^38^. The 27 strains were treated with 30 µM the compound library in 40 µl LB in black 384 well-plates with clear bottom (CatNo. 781097,

Greiner-Bio One, Germany). Precultures were grown overnight, diluted to an OD_600 nm_ of 0.1 and used to inoculate 384 well-plates with a Singer Rotor+ replicator (Singer Instruments, UK), resulting in ∼1:600 dilution. Plates were sealed with transparent breathable membranes (Breathe-easy^®^, Cat. No.: BEM-1, Diversified Biotech, USA) and incubated at 37°C and shaking at 800 rpm in our automated workstation. Growth (OD_600 nm_) and reporter activity (luminescence) were measured at regular 40 minutes intervals over 12 hours in the plate reader. The screen was done in biological duplicates.

The compound library used in this study was compiled from three commercial libraries purchased from MedChemExpress (USA): FDA-approved drug library (Cat. No.: HY-L022), Hulman Endogenous Metabolite Compound Library (Cat. No.: HY-L030), and Medicine Food Homology Compound Library (Cat. No.: HY-L055). The exact list of compounds is provided in Supplementary table 2.

### Data analysis pipeline

Data analysis was performed using R (version 4.2.2) and network visualization was done with Cytoscape 3.10.4. The data from compound-promoter combinations screen was analyzed in a similar fashion as previously described ^38^ with the following adjustments. The pipeline consists of three parts: i) quality control, ii) calculation of luminescence fold-change between treatment and untreated controls (*luxFC_treated_*_/*untreated*_), and iii) compound-promoter interactions (CPIs) calling. These steps are briefly described as follows:

i. Quality control: All growth curves were pre-processed with the R package DGrowthR ^81^. For each growth curve, the minimum of the 3^rd^ and 4^th^ OD_600 nm_ measurement was used as a baseline and its value was subtracted from all subsequent measurements. The initial two measurements were discarded due to technical noise. Afterwards, experiments where the maximum OD_600 nm_ was less than 20% of the maximum OD_600 nm_ recorded across all experiments were filtered-out (n=99). Compounds that inhibited the growth of > 50% of strains were discarded (n=10 compounds). Finally, non-replicable compound-promoter pairs, where the difference in OD_600 nm_ after 10 hours was greater than 0.5 were also not considered further. All excluded data is labelled in Supplementary table 3). In total, after quality control 59,842 unique compound-promoter pairs were considered for further analysis.
ii. Calculation of *luxFC_treated_*_/*untreated*_: The first two timepoints of each Lux curve were discarded, consistent with growth data pre-processing. The raw luminescence signal over time per well was summarized as the empirical area under the curve of luminescence over time until 10h of incubation (Lux*_auc_* _10ℎ_). Lux*_auc_* _10ℎ_ was normalized by the OD_600_ nm at the same timepoint (OD_10ℎ_) to account for the effect of population size on the magnitude of the luminescent signal:

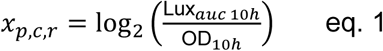

where *x_p_*_,*c*,*r*_ is the growth-normalized luminescence signal for a given promoter (*p*), compound (*c*), replicate (*r*). *x_p_*_,*c*,*r*_ was then median-centered for each promoter-replicate (*p*, *r*) combination, to make the magnitude of the experiments comparable:

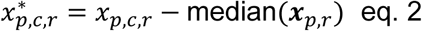

Compound specific activity was calculated by subtracting the average signal obtained from the EVC to the growth-normalized and median-centered luminescence signal *x*^∗^ of the same compound, in order to account for non-specific effects of the compound in the reporter system.

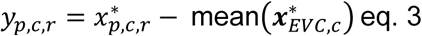

where *y_p_*_,*c*,*r*_ is the compound-specific effect on the reporter signal of a given promoter. For simplicity, *y_p_*_,*c*,*r*_ was renamed to *luxFC_treated_*_/*untreated*_, as it is a more intuitive name. In other words, *luxFC_treated_*_/*untreated*_ is numerically identical to *y_p_*_,*c*,*r*_.

iii. CPIs calling: For each promoter-replicate, DMSO experiments (***y****_p_*_,DMSO,*r*_ ∈ ℝ^128^) were normally distributed, with each promoter-replicate containing 128 DMSO wells. This was confirmed via a Lilliefors test for each promoter-replicate. Therefore, ***y****_p_*_,DMSO,*r*_ ∈ ℝ^128^ were used to parametrize a null normal distribution and calculate a respective z-score for each promoter-compound-replicate experiment:

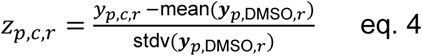

*z_p_*_,*c*,*r*_ were then used estimate p-values using a two-sided z-test:

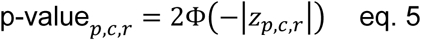

where Φ is the culmulative distribution function of the standard normal distribution. Finally, the largest of the two replicate p-values for each compound-promoter pair is retained (conservative measure), and all p-values for a given promoter are corrected for multiple testing via the Benjamini-Hochberg procedure. An adjusted p-value threshold of 0.05 was used to call significant CPIs. CPIs derived from *feoB*p with 695 compounds (distributed between two plates) were not considered, as a possible contamination of the second replicate of these plates could not be discarded (n=6 CPIs, Supplementary table 4). Nonetheless, the single-replicate was kept and used for the quantitative analysis.

### Rarefaction analysis

A rarefaction analysis via random subsampling of the chemical library was done to assess how the number of screened compounds affected detection of CPIs. For each subsample, the number of promoters that were differentially active in response to at least one compound within that subsample was quantified. Specifically, the complete set of compounds *C* ∈ 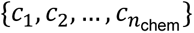 and promoters 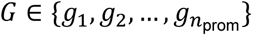, where *n*_chem_ = 2415 is the total number of screened compounds, and *n*_prom_ = 27 is the total number of screened promoters, was considered. For a given subset of compounds *C*^′^ ∈ *C* of size 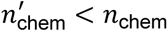, the total number of differentially expressed promoters was recorded. The value of 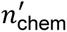 sequentially increases from 5 to *n*_chem_ − 1, with 100 random samplings performed at each subsample size. A similar procedure is followed for the promoter rarefaction analysis. For a random subsample of promoters *G*^′^ ∈ *G* of size 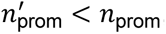, the number of compounds that affect at least one promoter in the subset is recorded. In this instance, the size of the subset of promoters was sequentially increased from 1 to *n*_prom_ − 1 in increments of 1.

### Concentration-resolved validation of CPIs

A validation set of 45 compounds was assembled, consisting of 40 compounds that exhibited at least one CPI in the primary screen, plus 5 compounds that showed no significant interactions (Figure S3a). Compounds were individually sourced from Biomol or Sigma-Aldrich-Merck (Supplementary table 17) and prepared as five-point, equally spaced dilution series. The maximum concentrations were determined based on each compound’s effect on *Salmonella* growth observed in the primary screen: compounds with no growth interference (n=37), mild effects (n=2), and considerable growth effects (n=6) were tested at maximum concentrations of 150 µM, 100 µM, and 50 µM, respectively.

Plate preparation and experimental procedures followed the same protocol as the primary screen. The effect of five concentrations of each compound was assessed using the same promoter library (n=26 promoters plus the empty vector control [EVC]), with all measurements performed in biological quadruplicates. For each well, growth (OD*_auc_* _10ℎ_) and reporter activity (Lux*_auc_* _10ℎ_) were quantified as area under the curve over 10 hours. Compound concentrations causing reduction >90% in maximum OD*_auc_* _10ℎ_ when compared to untreated controls were excluded from analysis. Data processing mirrored the primary screen protocol, with minor alterations. Growth normalization of the luminescence signal was performed as in the primaries screen, while median normalization (eq. 2) was done using only untreated samples:

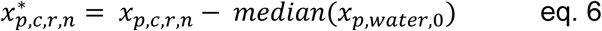

Similarly to the primary screen, compound specific activity was calculated by penalizing non-specific effects on EVC, in a concentration specific manner (*n*):

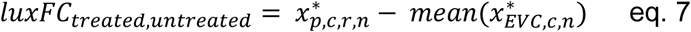

CPI calling was performed in the same manner as the primary screen, by using water-treated wells (n=48) as a null distribution. Significant CPIs are called when at least 2 of 4 biological replicates presented p-value < 0.01 and the absolute average of *luxFC_treated_*_,*untreated*_ was higher than 0.8. The experiments regarding the off-screen promoter STnc1480p were done using the same 45-compound validation set under identical experimental and data processing conditions.

To benchmark our results from the primary screen across different concentration thresholds, Receiver Operating Characteristic (ROC) and Precision-Recall (PRC) curves were generated using five concentration ranges from the validation set: ≤30, ≤60, ≤90, ≤120, and ≤150 µM. For each concentration range, CPIs from the primary screen (across 40 adjusted p-value thresholds ranging from 1 x 10^-150^ to 1.0) were compared against the concentration-resolved validation data, used as the reference standard. True positives, true negatives, false positives, and false negatives were determined, and area under the curve (AUC) values were calculated for both ROC and PRC curves for each concentration range.

We used Receiver Operating Characteristic (ROC) and Precision-Recall (PR) curves to assess the agreement between our primary screen and the findings made across different compound concentration thresholds. The CPIs identified at each concentration served as independent “ground truth” sets, while the adjusted p-values from the primary screen were used to rank each CPI. ROC and PR curves were then estimated across 40 p-value thresholds spanning from 1 x 10^-1^^50^ to 1.0.

### Individual promoter-reporter assays

Compound-mediated promoter activity for individual compounds and strains was used throughout this study to confirm specific findings from the primary screen and support molecular mechanism investigation. The assay was done similarly to the primary/secondary screens, using a single compound concentration, or more when indicated, and the respective vehicle. The experiments were done in black 96 well-plates with clear bottom (CatNo 655097, Greiner-Bio One, Germany) with 100 µl media - LB or N-minimal medium^27^ supplemented with 10 mM MgCl_2_, when indicated, starting with OD_600 nm_ of 0.005. The plates were incubated in a Infinite^®^ 200 PRO M Plex plate reader (Tecan Trading AG, Switzerland). at 37 °C with orbital shaking. Growth (OD_600 nm_) and reporter activity (luminescence) were measured at regular 30 minute intervals over 16 hours using the software i-control™ 2.0 (Tecan Trading AG, Switzerland). Data was analyzed R (version 4.2.2) as for the primary screen, using luminescence signal of the vehicle for median centering (similar to eq. 6).

A similar approach was used to compare basal promoter expression (compound free) changes across different strain backgrounds, where growth-normalized luminescence of a given strain was mean-centered by the wild type (eq. 8).

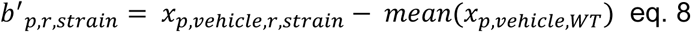

Finally, *b*^′^*_p_*_,*ve*ℎ*icle*,*r*,*s*_was then penalized for potential interference with the reporter system by using the EVC:

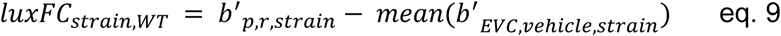

### Chemical representation and physicochemical properties calculation

Morgan fingerprints and physicochemical properties of all compounds in our chemical library were assembled using the RDKit v24.09.06 python module. First, all salts were removed from SMILES strings using the RemoveSalts() function with default parameters. Binary Morgan Fingerprints are gathered using the GetMorganGenerator() function with the parameter radius=2. A 2048-bit vector is obtained for each compound. Additionally, a set of 98 physicochemical properties were calculated as previously described ^82^.

Data from the complete DrugBank database was downloaded in June 2025 ^45^. Only compounds with a molecular weight ≤ 1000 Daltons were considered for analysis. Morgan fingerprints and physicochemical properties were assembled/quantified as described above.

Molecular dissimilarity based on Morgan fingerprints was estimated using the Jaccard distance. For a given pair of compounds *i* and *j*, the Jaccard distance (*δ_i_*_,*j*_) of the corresponding Morgan fingerprints 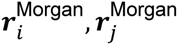 is estimated as:

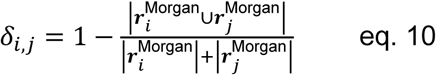

Following previous convention, two compounds were considered “dissimilar” if their Jaccard distance is greater than 0.7 ^83^.

### RNA extraction, cDNA library preparation and RNA sequencing

Overnight cultures of *Salmonella* wild type, Δ*marA* and Δ*slyA* were diluted to OD_600 nm_ ∼0.01 in LB, grown until OD_600 nm_ ∼0.1 and dantrolene (100 µM) or DMSO were added for 1 hour. Cultures were mixed with 0.1 volumes of stop mix (95% EtOH and 5% phenol, v/v), centrifuged for 10 min at 4000 x g and 4 °C, and the supernatant was discarded. Pellets were snap-frozen in liquid nitrogen and stored at –80 °C. Frozen pellets were thawed on ice and resuspended in 600 µl Tris-EDTA buffer (pH 8.0) containing 0.5 mg/ml lysozyme and 1% SDS. Cells were lysed upon 2 min incubation at 64 °C and total RNA extraction was conducted using the hot phenol method as described previously ^84^. After RNA isolation, residual DNA was removed by treatment with DNase I (Thermo Scientific) according to the manufacturer’s instructions.

Ribosomal RNA depletion and cDNA library preparation of RNA samples from DMSO- and dantrolene-treated *Salmonella* WT, Δ*marA* or Δ*slyA* were performed by the Core Unit SysMed Würzburg, Germany. A DNase digest (Thermo Fisher Scientific) was performed followed by depletion of ribosomal RNA molecules using RiboCop META (Cat. No. 125) rRNA depletion kit (Lexogen). The ribo-depleted RNA samples were first fragmented using ultrasound (2 pulses of 30 s at 4°C). Then, an oligonucleotide adapter was ligated to the 3’ end of the RNA molecules. First-strand cDNA synthesis was performed using M-MLV reverse transcriptase with the 3’ adapter as primer. After purification, the 5’ Illumina TruSeq sequencing adapter was ligated to the 3’ end of the antisense cDNA. The resulting cDNA was PCR-amplified using a high-fidelity DNA polymerase and the barcoded TruSeq-libraries were pooled in approximately equimolar amounts. Sequencing of pooled libraries, spiked with PhiX control library, was performed at 10 million reads per sample in single-ended mode with 100 cycles on the NextSeq 2000 platform (Illumina). Demultiplexed FASTQ files were generated with bcl-convert v4.3.6 (Illumina).

Genomic annotations for *Salmonella enterica* subsp. *enterica* serovar Typhimurium str. 14028S were gathered under GenBank accession CP001363.1. RNA-seq fastq file processing was performed using the singularity profile of the nf-core/rnaseq pipeline v3.16.0 (10.5281/zenodo.1400710) in order to obtain transcript quantifications. Differential expression analysis was performed using DESeq2 v1.48.1. An adjusted p-value threshold of 0.05 was used to determine significant differential expression. Shrinked log fold change estimation is performed with the lfcShrink() function with type=”apeglm” ^85^.

### Reverse transcription quantitative polymerase chain reaction (RT-qPCR)

Overnight cultures of *Salmonella* wild type were diluted to OD_600 nm_ ∼0.01 in LB and dantrolene (100 µM) or DMSO were added either immediately and for the duration of 3 hours, or when OD_600 nm_ reached 0.1 (about 2 h) for the duration of 1h. Quantification of mRNA levels of *virK* were performed for DNase I-digested RNA samples after DNA extraction as described above using the Maxima H Minus First Strand cDNA Synthesis Kit (Thermo Fisher Scientific). One microgram of RNA was denatured, snap-cooled on ice, and reactions were started with or without Maxima H Minus™ RT using the following cycling conditions: 25 °C, 10 min; 50 °C, 30 min; 85 °C, 5 min. RT-PCR reactions were prepared using the kit Takyon® No Rox SYBR Core Kit blue dTTP (Eurogentec, Belgium), following manufacturer’s instructions. PCR was conducted using gene-specific oligonucleotides for *virK*, and the reference *rpoS* (supplementary table 14) in a CFX Opus 384 PCR system (Bio-Rad), and analyzed in the software Bio-Rad CFX Maestro 2.3 (Bio-Rad). Experiments were conducted in technical duplicates of three biological replicates for DMSO- or dantrolene-treated samples.

### *In silico* structural analysis of SlyA binding to dantrolene and triflusal

A strategy based on structure-based modelling, induced-fit molecular docking, molecular dynamics simulations, and MM/GBSA binding energy calculations was followed to characterize the interaction of dantrolene and triflusal with *Salmonella enterica* SlyA *in silico*. Starting from experimentally resolved SlyA conformations, molecular docking was used to identify potential ligand-binding pockets and evaluate ligand-induced conformational changes, and finally, the stability and energetics of protein-ligand complexes over time were assessed by molecular dynamics simulations. Protein preparation, pocket identification, molecular docking, and molecular dynamics simulations were performed using the Schrödinger Drug Discovery Suite (version 2024.3). A complementary section containing detailed results to support the modelling strategy is provided as supplementary information.

#### Protein structure preparation and molecular docking

Structural models of *Salmonella* enterica SlyA were generated from experimentally resolved structures retrieved from the Protein Data Bank (PDB). Missing loops and unresolved side-chain atoms were reconstructed to obtain complete models suitable for computational analyses. To account for conformational differences associated with DNA and ligand binding, the salicylate-bound SlyA structure (PDB 3DEU) was aligned against the DNA-bound SlyA structure (PDB 3QF5), generating comparable structural models for ligand-binding analyses ^60^. Protein structures were prepared using the Protein Preparation Wizard, including assignment of protonation states, addition of hydrogen atoms, reconstruction of missing side chains, and optimization of hydrogen-bonding networks. Ionization states were selected according to the most probable assignments predicted by the software at physiological pH. All structures were subsequently minimized using the steepest descent algorithm with a convergence cutoff of 0.5 Å.

Potential ligand-binding pockets were identified and evaluated using SiteMap based on predicted pocket geometry, volume, and druggability scores. Dantrolene and triflusal were drawn and prepared using LigPrep to generate three-dimensional conformations, assign protonation states at pH 7.4, and calculate partial atomic charges using the OPLS4 force field ^86,87^. Whenever alternative protonation or tautomeric states were predicted, all plausible ligand forms were considered for docking analyses. Ligand docking was performed using Glide (version 7.7) in induced-fit docking mode with standard precision (SP), followed by local side-chain refinement using Prime ^88,89^. Docking grids were centered either on the predicted SiteMap binding pockets or on the coordinates of co-crystallized ligands when applicable, using a 12 Å search region from the pocket centroid ^90^. The induced-fit protocol enabled simultaneous sampling of ligand flexibility and local protein side-chain rearrangements to account for conformational adaptation upon ligand binding.

#### Molecular dynamics simulations

The stability and dynamics of the resulting protein–ligand complexes were evaluated using all-atom molecular dynamics (MD) simulations in explicit solvent conditions using the Desmond simulation package with the OPLS4 force field ^87^. Protein–ligand complexes were solvated using the TIP3P water model and neutralized with counterions ^91^. Systems were placed in cubic periodic boundary boxes with a minimum distance of 10 Å between the protein and box edges. Simulations were conducted with a 1 fs integration time step. Short-range electrostatic interactions were calculated using a 9.0 Å cutoff, while long-range electrostatics were treated using the Particle Mesh Ewald (PME) method ^92^. Energy minimization and system relaxation were performed using a hybrid protocol combining Steepest Descent and limited-memory Broyden–Fletcher–Goldfarb–Shanno (L-BFGS) algorithms, following the default Desmond relaxation workflow. Equilibration was carried out under the NPT ensemble for 5 ns using the Berendsen thermostat and barostat ^93^. Production simulations were maintained at 310 K using the Nosé–Hoover thermostat and at 1 atm using the Martyna–Tobias–Klein barostat ^94,95^. Production runs were performed for at least 300 ns, with trajectory frames saved every 1 ns. Ten independent replicas were generated for each compound, resulting in a cumulative simulation time of ∼3 μs per ligand. For hit compounds, 5 independent simulations of 300–500 ns were performed. Trajectory and interaction data are publicly available in the Zenodo repository (DOIs: 10.5281/zenodo.15582334 and 10.5281/zenodo.17858054) ^96^.

Representative structures were selected based on stabilization of root-mean-square deviation (RMSD) values after equilibration. RMSD, RMSF, and interaction analyses were performed using the Maestro Simulation Interaction Diagram tool (Schrödinger LLC). Structural visualization was carried out in PyMOL v2.5.4–3.1, and data visualization was performed using Python 3.7 with matplotlib, seaborn v0.12.2, and GraphPad Prism v10.1 ^97^.

Finally, binding free energies of protein-ligand complexes were estimated using the molecular mechanics/generalized Born surface area (MM/GBSA) approach. Every tenth frame from the MD trajectories was extracted and used as input for MM/GBSA calculations performed with the thermal_mmgbsa.py script from the Schrödinger suite using Prime 14 ^98^. Calculated binding free energies (ΔG) were additionally normalized by the number of ligand heavy atoms (HAC) to estimate ligand efficiency (LN), according to the equation:

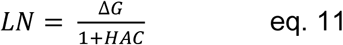

### Polymyxin-B sensitivity assay

The assay was performed as previously described ^99^ with minor modifications. Overnight cultures of *Salmonella* wild type, Δ*slyA*, and *slyA* W34A were grown at 37°C in N-minimal medium (NMM, pH 7.4) supplemented with 10 mM MgCl₂. Cells were washed three times with MgCl_2_-free NMM and diluted 1:100 into NMM containing 10 µM MgCl₂, either with dantrolene (100 µM), triflusal (100 µM), or the corresponding DMSO vehicle control (1.38% and 0.5% DMSO for dantrolene and triflusal, respectively). 2 mL cultures were incubated for 3 h at 37°C with shaking at 200 rpm. Approximately 1 × 10⁷ cells from each sample were then inoculated into LB medium containing 5 µg/mL polymyxin-B (Sigma) and incubated for 1 h at 37°C with shaking at 200 rpm to assess viability. Serial dilutions were prepared in PBS and plated on LB agar in technical duplicates. Percentage survival was calculated based on colony forming units (CFU) counting in relation to the initial inoculum (t = 0 h).

### SPI-2 type III secretion system activity assay

The secretion of the effector protein SseF was used to quantify the activity of the SPI-2 type III secretion system (T3SS-2). The SseF-HiBiT secretion assay was done as previously described ^62,100^. *Salmonella enterica* serovar Typhimurium NCTC 12023 was grown overnight in 2 mL LB. Cultures were washed twice with 1 mL PCN medium ^101,102^, pH 5.8, by centrifugation at 6000 × g for 2 min. Cultures were back-diluted to an OD_600_ of 0.1 in 1 mL PCN medium, pH 5.8, supplemented with triflusal at desired concentration or DMSO 1% (v/v), then incubated for 7 hours at 37°C with shaking at 180 rpm. Samples of 100 µL were transferred into a transparent 96-well plate for OD_600_ measurement. Subsequently, samples of 25 µL were transferred into a white 384-well plate. For the quantification of SseF-HiBiT, Nano-Glo® HiBiT Extracellular Detection System (Promega) was used. First, the HiBiT substrate (1:50 dilution) and LgBiT (1:100 dilution) were added into the HiBiT extracellular buffer. Aliquots of 25 µL of the mixture were then added to each well containing the samples. The plate was then incubated for 10 minutes in the dark before luminescence was measured using a Tecan Spark® Multimode Microplate Reader. Dose-response curves and IC_50_ values were calculated using GraphPad Prism 10.1.1.

### Cell culture and infection assay

The infection of HeLa-S3 cells was carried out as previously described ^103^, with minor adjustments. Two days prior to infection, 2×10⁵ cells were seeded in 2 ml of medium in six-well plates. One colony of *Salmonella* SL1344 was inoculated into 3 ml of LB liquid medium and grown overnight at 37 °C and 200 rpm. An overnight culture of *Salmonella* SL1344 was diluted 1:20 in LB medium and grown to an OD_600nm_ of 2.0, harvested by centrifugation for 2 minutes at 12,000 rpm at room temperature, and then resuspended in DMEM medium. Infections were performed at a multiplicity of infection (MOI) of 20 by adding the bacterial suspension to each well. To synchronize the infection, the six-well plates were centrifuged for 10 minutes at 250 x *g* at room temperature, followed by a 30-min incubation in a 5% CO₂ incubator. To kill extracellular bacteria, the medium was aspirated and replaced with DMEM containing gentamicin at a final concentration of 100 µg/ml. After a 30 min incubation at 37 °C, the medium was aspirated and replaced with DMEM containing gentamicin at a final concentration of 10 µg/ml. Dantrolene (60 µM), triflusal (100 µM) or the DMSO controls (0.8% or 0.2% as vehicle controls of dantrolene and triflusal, respectively) were added at this stage of the infection protocol. Eight hours after gentamicin was first added to the cells, the infected HeLa-S3 cells were solubilized using DPBS containing 0.1% Triton. For plating of colony forming units (CFUs), the wells were washed twice with 1x DPBS (Gibco) and 2 ml of 0.1% Triton-x DPBS were added to selectively lyse the host cells. After 5 min at room temperature, the cells were scratched off the bottom of the wells using a cell scraper, resuspended using a 1 ml tip and serially diluted before plating on LB plates for overnight growth at 37 °C. The number of CFUs was counted for at least three biological replicates.

### Compound cytotoxicity assay

Dantrolene and triflusal cytotoxicity on HeLa-S3 cells was assessed by measuring lactate dehydrogenase (LDH) release using the CytoTox 96® (Promega) assay. Briefly, 5,000 HeLa-S3 cells were dispensed into a 96-well plate the day before the experiment. The cells were incubated in a 5% CO_2_ humidified atmosphere at 37°C. The supernatant was then aspirated and 100 µL of DMEM (Gibco) supplemented with 10% FCS long with increasing concentrations (30 to 150 µM) of dantrolene or triflusal. Untreated or vehicle DMSO (0.8% and 2%) were used as controls. Treated wells were incubated at 37°C with 5% CO_2_ for 24 h. The relative proportion of dead cells was determined by lysing untreated HeLa-S3 cells with 1x Lysis Solution (Promega), according to the manufacturer’s instructions. Formazan formation, that is proportional to the number of lysed cells, was measured by monitoring the absorbance value at 490 nm using an Infinite^®^ 200 PRO M Plex plate reader (Tecan Trading AG, Switzerland). The average value of the culture medium background was subtracted from all experimental well values. At least two biological replicates were analyzed, with each replicate comprising three technical replicates.

### Minimal Inhibitory Concentration (MIC) determination

*Salmonella* overnight cultures were back-diluted as described for the primary screen, and used to inoculate 384-well microtiter plates (cat. no. 781162, Greiner Bio-One) containing 40 µL of LB medium with increasing concentrations of ciprofloxacin or chloramphenicol. For each antibiotic, a linear dilution series consisting of 11 equal steps was prepared, enabling high-resolution quantification of antimicrobial activity. The maximum concentration tested was set at 4x the previously determined strain-specific MIC for ciprofloxacin and chloramphenicol. Bacterial fitness for each antibiotic concentrations was calculated by normalizing the area under the growth curve (OD_AUC 8h_) to that of control wells without no antibiotic. MIC curves were generated by fitting a three-parameter logistic model to the antibiotic dose-response data using the drc package in R, as previously described ^104^.

### Ciprofloxacin killing assays

*Salmonella* overnight cultures were diluted 1:100 in LB and grown at 37 °C and 200 rpm until mid-exponential phase (OD_600 nm_ ∼0.4). Cells were diluted at 1 × 10⁷ CFU/mL in the presence of antibiotic ranges based on previously-determined inhibitory concentrations of ciprofloxacin (2 - 10x MIC) with or without the addition of dicyclomine (100 µM) or azelastine (100 µM). Killing was performed for 5 h at 37°C with shaking at 200 rpm. Afterwards, cultures were diluted in PBS and plated on LB agar in technical duplicates. Absolute colony forming units (CFU) counting of samples were obtained for three independent experiments.

### LASSO logistic regression model

Lasso logistic regression models were trained to predict whether a compound can trigger *ramA* up-regulation based on a representation of its molecular structure. Morgan fingerprints or the set of tabular chemical descriptor features (Supplementary table 10) were used as input molecular features. Prior to modelling, input feature matrices were pre-processed to remove correlated and non-informative features. In the case of the Morgan fingerprints, features that were present or absent in all compounds were removed, as well as features that were not present in at least 10 compounds. Additionally, correlated features with a hamming distance < 0.01 were also excluded. In the end, 1054 features remained. This resulted in an input feature matrix 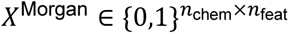, where *n*_chem_ is the number of compounds and *n*_feat_=1054 is the number of features. A similar procedure was employed for chemical descriptor features, ensuring a minimum feature variance of 0.1 and removing features with a Pearson correlation coefficient ≥ 0.9. A total of 51 features were retained for modelling, resulting in an input feature matrix 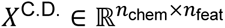. Compounds were assigned into training (80%), validation (10%) and testing (10%) sets following the scaffold splitting procedure ^105^. During model training, chemical descriptor features were mean-centered and scaled. Validation and test sets were transformed using the scaling parameters estimated for the training set.

A LASSO logistic regression model was trained to estimate the probability of a compound being a *ramA* inducer, using the corresponding molecular representation.

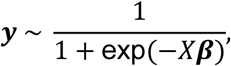

where 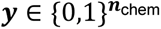 is a binary vector indicating whether a compound is an inducer or not. The optimal values for the model parameters (***β***) are estimated via Maximum Likelihood Estimation (MLE). To find the most parsimonious model, the LASSO regularization ^106^ was used, which adds a penalty term to the likelihood function, encouraging a sparse coefficient vector. In this way, the LASSO becomes a form of feature selection, prioritizing the contribution of the most informative features to predict the outcome. The complete loss function is defined as:

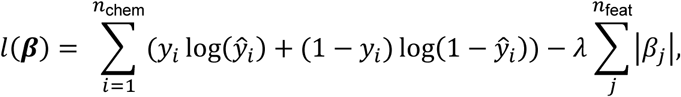

where *y_i_* indicates whether compound *i* is a hit, *ŷ_i_* is the predicted probability of *i* being a hit, and λ is a regularization parameter controlling the strength of the penalty term. The R package glmnet v4.1-9 ^107^ with type.measure=”auc” was used optimize model parameters by maximizing *l*(***β***). The optimal value for λ is found via 10-fold cross-validation on the training set. The lambda value that minimized the loss function *λ*_min_and the value that resulted in a loss value within one standard error of the minimum *λ*_1se_ were compared for generalization performance in the validation set. The area under the precision-recall curve (PR-AUC) is estimated for the validation set. The probability threshold that maximized f1-score was also determined on the validation set and was used to binarize prediction scores (model scores, threshold = 0.44) into predicted labels. For both feature-sets, *λ*_1se_ performed best. Finally, models trained on Morgan fingerprints and chemical descriptors were compared on the testing set.

DrugBank compounds were featurized to match features used for model training. The model score for each compound to be a *ramA* inducer is then binarized using the threshold found in the validation set (0.44).

